# Convection-enhanced delivery of dexamethasone in glioma suppresses myeloid inflammation while avoiding systemic toxicities

**DOI:** 10.1101/2025.09.24.677899

**Authors:** Nathaniel W. Rolfe, Nicholas B. Dadario, Liang Lei, Anthony Tang, Misha Amini, Damian E. Teasley, Nkechime Ifediora, Peter Chabot, Nathan J. Winans, Nina Yoh, Julia Furnari, Corina Kotidis, Clara H. Stucke, Nivia M. Urena, Yanping Sun, Abby Brand, Ashwin Viswanathan, Pavan Upadhyayula, Michael G. Argenziano, Colin P. Sperring, Nadine Khoury, Nelson Humala, Justin Neira, Peter A. Sims, Brian J. Gill, Peter Canoll, Jeffrey N. Bruce

**Author notes:** Correspondence: Jeffrey N. Bruce, Department of Neurological Surgery, Columbia University Medical Center, 710 W. 168th St., New York, NY 10032. Nathaniel W. Rolfe, Nicholas B. Dadario, Liang Lei, and Anthony Tang are co-first authors.

## Abstract

Dexamethasone is widely used to control cerebral edema and inflammation in glioblastoma, but its benefits are limited by systemic toxicities and adverse prognostic associations. We evaluated local administration of dexamethasone via convection-enhanced delivery (CED) to maximize intratumoral anti-inflammatory effects by increasing local corticosteroid exposure while minimizing systemic exposure. In two glioma mouse models, continuous intraparenchymal infusion of dexamethasone was well tolerated with no adverse effects. Pharmacokinetic analyses supported preferential intratumoral distribution and reduced systemic exposure with CED compared with systemic dosing. Single-nucleus RNA sequencing (snRNA-seq) and immunohistochemistry showed attenuation of glioma-associated inflammation with downregulation of reactive microglial/macrophage programs and reduced tumor-infiltrating myeloid cells with a morphology consistent with a less activated state. Experiments in human induced pluripotent stem cell (iPSC)–derived microglia confirmed that dexamethasone directly suppresses inflammatory gene expression, indicating a conserved mechanism across species. This inflammatory suppression was recapitulated in both immortalized microglial (HMC3) and macrophage (THP1) cell lines. These findings suggest that localized dexamethasone delivered by CED reprograms the glioma immune microenvironment and achieves control of inflammation without the systemic adverse effects associated with standard systemic dexamethasone therapy. This clinically translatable strategy may improve symptom management and provide a platform for integrating local immunomodulation with future glioblastoma therapies.

## Introduction

Glioblastoma (GBM) is the most common and aggressive primary brain malignancy and is associated with a dismal clinical prognosis (1). Despite standard of care treatment consisting of surgical resection, temozolomide, and radiotherapy, median survival for glioblastoma is around 15 months after diagnosis, a consequence of the nearly inevitable recurrence of the disease (2).

Cerebral edema and tumor-associated inflammation are among the chief clinical problems contributing to neurological decline and reduced quality of life in glioblastoma patients. Since the 1960s, these problems have primarily been managed with dexamethasone (3). Patients are frequently on dexamethasone therapy for prolonged periods of time, often until death (4, 5). As a result, Cushing syndrome is common (4, 6) with irritability, weight gain, muscle atrophy, plethora, moon facies, ecchymoses, striae, hyperglycemia, and systemic immunosuppression (7). These adverse effects likely contribute to the observed negative correlation between dexamethasone dose and overall survival in glioblastoma patients (8, 9).

Dexamethasone exerts its effects through both genomic and nongenomic mechanisms (10–12), and its pleiotropic effects are responsible for both its potent edema-reducing effects and its harmful side effects (13). Improvements in cerebral edema following dexamethasone therapy have been linked to its suppression of vascular permeability factors, including VEGF and IL-1β, as well as upregulation of various tight-junction proteins in endothelial cells (14–18). Dexamethasone has been shown to attenuate microglial activation to inflammatory stimuli both in vitro and in vivo and decrease nitric oxide and cytokine production by activated microglia (19–22). In vivo glioblastoma studies have demonstrated the immunosuppressive effects of dexamethasone on T-cells (23, 24), but concomitant effects on the chronic inflammatory milieu remain poorly defined.

The diffuse nature of gliomas means that neoplastic cells grow and intermingle with non-neoplastic cells in the tumor microenvironment. The non-neoplastic component of the glioma microenvironment primarily consists of tumor-associated microglia and macrophages, which comprise up to 30% of tumor bulk in primary glioblastoma (25, 26). This chronic inflammatory environment promotes tumor progression and is therefore a putative therapeutic target (27–30). Broad inhibition of tumor-associated microglia and macrophages with CSF-1R inhibitors and targeted suppression of IL-1β have both been shown to prolong survival in glioma-bearing mice (28, 29, 31). Unfortunately, an initial clinical trial of a CSF-1R inhibitor was unsuccessful (32), and IL-1β inhibition has limited clinical precedent in glioblastoma. Dexamethasone remains the most clinically relevant and widely used anti-inflammatory drug in glioblastoma patients.

CED is a high-precision local drug infusion methodology in which drugs are delivered at high concentrations directly into the tumor and peritumoral brain through a surgically implanted catheter attached to a microinfusion pump (33–35). Compared to systemic drug delivery, CED achieves higher drug concentrations in the brain while minimizing systemic toxicities (36, 37). Initial studies of CED used an external pump and catheter, which prevented continuous or repeated drug infusion without the danger of infection or the risks of repeat surgery (38, 39).

While this limited the efficacy of early CED trials, the recent successful clinical trial use of subcutaneously implantable, refillable pumps for repeated and prolonged drug infusions has expanded CED’s clinical advantage (40, 41). By facilitating long-term delivery and repeat dosing, a chronic-CED strategy allows the use of multiple drugs, in series or parallel, that would not be practical with systemic delivery (42). The advantages of chronic CED are well-suited for dexamethasone, which can exert its effects in the peritumoral environment over a prolonged period while avoiding unwanted systemic toxicities.

We utilized a syngeneic glioma mouse model to compare systemic toxicities and the effects on the tumor-associated inflammatory microenvironment when dexamethasone is delivered locally via CED versus systemically via daily i.p. injection. We first demonstrate that CED of dexamethasone is well tolerated in glioma-bearing mice and allows for preferential accumulation within the brain parenchyma compared to systemic administration. Additionally, we show that CED of dexamethasone avoids the systemic toxicities that accompany systemic dosing. We also observe a reduction in inflammatory transcriptional signatures and histologic microgliosis with CED of dexamethasone. Finally, we show that the anti-inflammatory effects of CED-dexamethasone are recapitulated in vitro using human induced pluripotent stem cell-derived (iPSC) microglia.

## Results

### Convection-enhanced delivery of dexamethasone is well tolerated in glioma-bearing mice

Local delivery of a high dose of dexamethasone (0.24 mg/day) has been shown to prolong survival in a leporine metastatic carcinoma model (43). Therefore, we performed a survival study to determine the tolerability and potential benefit of locally delivered dexamethasone in a syngeneic glioma model. Mice were randomized to receive either a phosphate buffer solution (PBS) control or PBS solution containing 100 ng/μl of dexamethasone via a seven-day Alzet pump (1007D). An intra-pump concentration of 100 ng/μl of dexamethasone was chosen as it was below the level affecting tumor cell viability in our PDGFA-driven syngeneic murine glioma cell line and GL261 cells (Supplementary Figure 1A). This was important as we wanted to explore dexamethasone’s effects on the inflammatory microenvironment rather than its direct cytotoxic effects. Additionally, mice that received an intra-pump concentration of 200 ng/μl of dexamethasone exhibited nearly 10% weight loss one day after pump implantation (Supplementary Figure 1B), suggesting that 100 ng/μl is near the murine maximally tolerated dose when delivered via a seven-day Alzet pump. Mice receiving systemic dexamethasone received daily i.p. injections of 10 mg/kg of dexamethasone, a dose found tolerable in previous survival studies (8).

All mice demonstrated adequate intraparenchymal distribution of the drug or vehicle as assessed by co-infused gadolinium (1% Omniscan), and a consistent volume of distribution was observed between groups (Figure 1A). CED-dexamethasone treatment for seven days was well-tolerated without significant changes in weight compared to control mice or mice receiving systemic dexamethasone (Figure 1B) or outward signs of distress.

**Figure 1:**
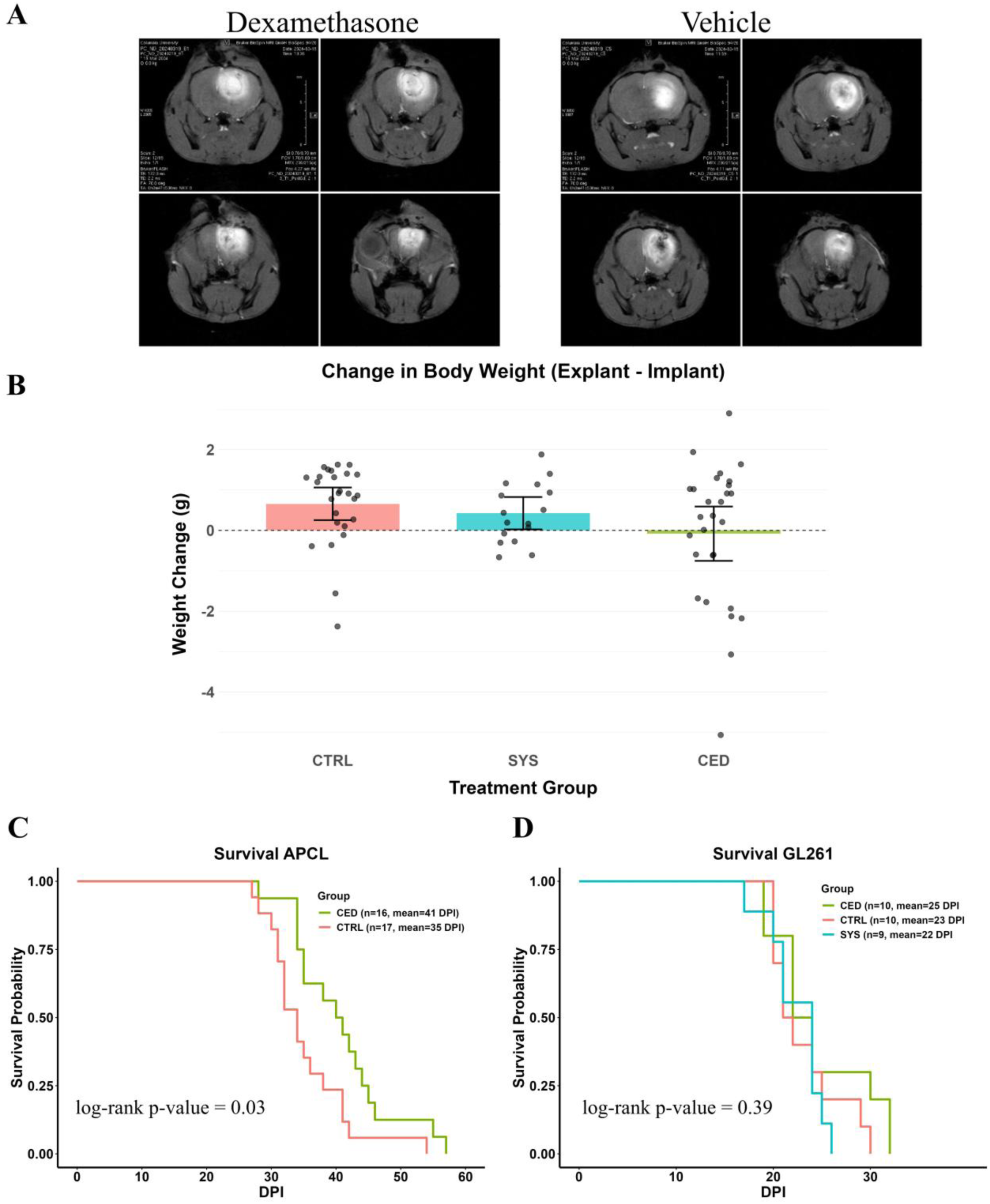
**A** – Representative MRI images from mice demonstrating the volume of gadolinium distribution between control and CED dexamethasone treatment conditions. **B** – The majority of mice gained or lost a negligible amount of weight in our three treatment conditions. Mice receiving PBS (vehicle control) gained 0.7 ± 1.0 g (mean ± SD), mice receiving systemic dexamethasone gained 0.4 ± 0.8 g, and mice receiving CED-dexamethasone lost 0.1 ± 1.7 g. There was no significant difference in weight change between treatment groups (Kruskal-Wallis, p = 0.15). **C** – Survival curves of vehicle- or CED-dexamethasone-treated mice with APCL tumors. Mean survival for control mice was 35 days while median survival for CED-dexamethasone-treated mice was 41 days (log-rank p = 0.03). **D** – Survival curves of vehicle-, CED-dexamethasone, and systemic dexamethasone treated mice with GL261 tumors. Mean survival for control mice was 23 days, mean survival for systemically treated mice was 22 days, and mean survival for CED-dexamethasone treated mice was 25 days (log-rank p = 0.39).

Kaplan-Meier survival analysis demonstrated a modest but statistically significant survival advantage in our APCL model for mice treated with CED of dexamethasone compared to vehicle-treated control mice (mean survival 41 DPI vs 35 DPI, respectively; log-rank p = 0.03) (Figure 1C). This survival effect did not generalize to the GL261 model, where mice receiving CED-dexamethasone, systemic dexamethasone, and vehicle control did not demonstrate a statistically significant change in overall survival (mean survival 25 DPI, 22 DPI, and 23 DPI, respectively; log-rank p=0.39) (Figure 1D).

### CED of dexamethasone improves drug delivery to the brain while minimizing systemic exposure

Using high-performance liquid chromatography-tandem mass spectrometry (HPLC-MS/MS), we quantified dexamethasone concentrations in tissue and plasma under various delivery conditions in both murine and human specimens.

We first showed that dexamethasone is stable at 37°C for up to 10 days in the Alzet pump. (Supplementary Table 1). After five days of local dexamethasone delivery in mice, the average concentration of dexamethasone in the tumor-bearing quadrant was 196.2 ng/g (Figure 2A). The concentration of dexamethasone in the liver of CED-treated mice was 6.1 ng/g, which likely results from dexamethasone efflux from the brain (Figure 2A). The dexamethasone concentrations in the liver, brain, and serum of mice were measured one hour after i.p. injection of 10 mg/kg of dexamethasone (Figure 2B). When delivered systemically, dexamethasone preferentially partitions in the liver (10,998.9 ng/g versus 2,410.9 ng/mL of serum), with a relatively lower concentration in the brain (124.8 ng/g). The partitioning coefficient between brain and liver parenchyma of K_brain,liver,ip_ = 0.011 ± 0.010, whereas local delivery allows for preferential partitioning in the brain with a K_brain,liver,ced_ = 29.2 (Figure 2C).

**Figure 2:**
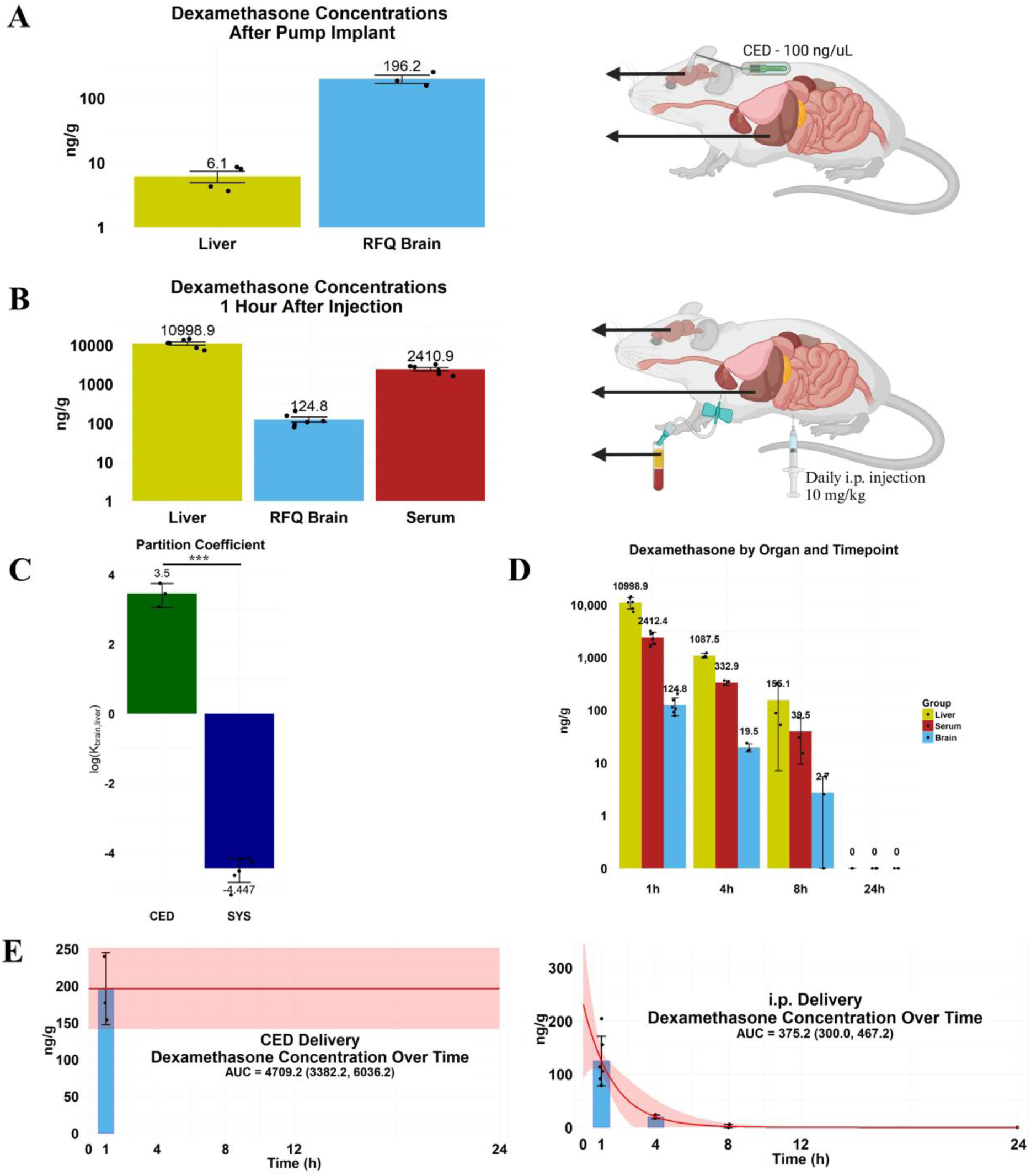
Convection-enhanced delivery of dexamethasone leads to preferential and sustained drug exposure in the tumor-bearing brain with minimal peripheral distribution compared to systemic administration. **A** – Dexamethasone concentrations in the tumor-bearing quadrant (196.2 ± 56.4 ng/g) and liver (6.7 ± 3.1 ng/g) in mice treated with CED of dexamethasone. Values were obtained after 5 days of treatment with an Alzet osmotic pump (1007D). **B** – Dexamethasone concentration in the tumor-bearing quadrant (124.8 ± 93.4 ng/g), liver (10,999 ± 5,486 ng/g), and serum (2,411 ± 1,248 ng/g) of mice one hour after i.p. injection of 10 mg/kg of dexamethasone. **C** – The partitioning of dexamethasone between the brain and liver depends on the route of administration. CED of dexamethasone allowed for preferential partitioning of the drug in the brain. Intraperitoneal injection led to a partitioning coefficient of K_brain,liver_ = 0.011 ± 0.010. CED led to a partitioning coefficient of K_brain,liver_ = 29.2 ± 15.8. **D** – Dexamethasone concentrations in the brain, liver, and serum were determined 1, 4, 8, and 24 hours post-i.p. injection of 10 mg/kg dexamethasone. **E** – Areas under the curve were calculated for once daily 10 mg/kg dexamethasone i.p. and for CED of dexamethasone (100 ng/μl). Assuming CED leads to a steady state concentration over 24 hours, we found that CED delivery gave an AUC (95% CI) of 4,709 (3382, 6036) (ng/g)·h, whereas systemic delivery had an AUC of 375 (300, 467) (ng/g)·h. (Values are reported as mean ± SE)

Additionally, we collected tissue at 1-, 4-, 8-, and 24-hours post i.p. dexamethasone injection and observed a first-order rate of elimination in both the liver, brain, and serum consistent with a half-life of approximately 1.1 hours in all three compartments (Figure 2D). We estimated the area under the curve (AUC) for the dexamethasone concentration over time when given via CED and via daily i.p. dosing. The AUC with CED of dexamethasone was 10-fold higher compared to systemic delivery (4,709 ± 1,327 vs 375 ± 75 (ng/g)·h, respectively; Figure 2E).

We also quantified dexamethasone partitioning in human brains and plasma from intraoperative GBM samples (Supplementary Figure 2A). Contrast-enhancing tumor samples had a concentration of dexamethasone of 73 ± 20% of the corresponding plasma concentrations. Non-enhancing samples, however, had a dexamethasone concentration of 20 ± 10% of the plasma concentration. In our murine studies, mice that received systemic dexamethasone had a dexamethasone concentration in the tumor-bearing quadrant of 6 ± 2% of the plasma concentration.

### CED of dexamethasone avoids side effects that accompany systemic administration of dexamethasone

Next, we demonstrated that CED of dexamethasone was a viable strategy to avoid systemic side effects. Systemically treated mice were given 10 mg/kg/day of dexamethasone via i.p. injection, while locally treated mice were given dexamethasone via an Alzet pump (1007D) at an intra-pump concentration of 100 ng/μl.

Systemically delivered dexamethasone led to several measurable physiological side effects in our mice. Blood glucose levels, taken 24 hours post i.p. dexamethasone dosing (a trough level), were decreased from their pretreatment baseline in systemically treated mice (Figure 3A), and a similar decrease was still seen on day seven of treatment (Figure 3B). When dexamethasone was given via CED, no change in blood glucose level was seen (Figure 3A, B).

**Figure 3:**
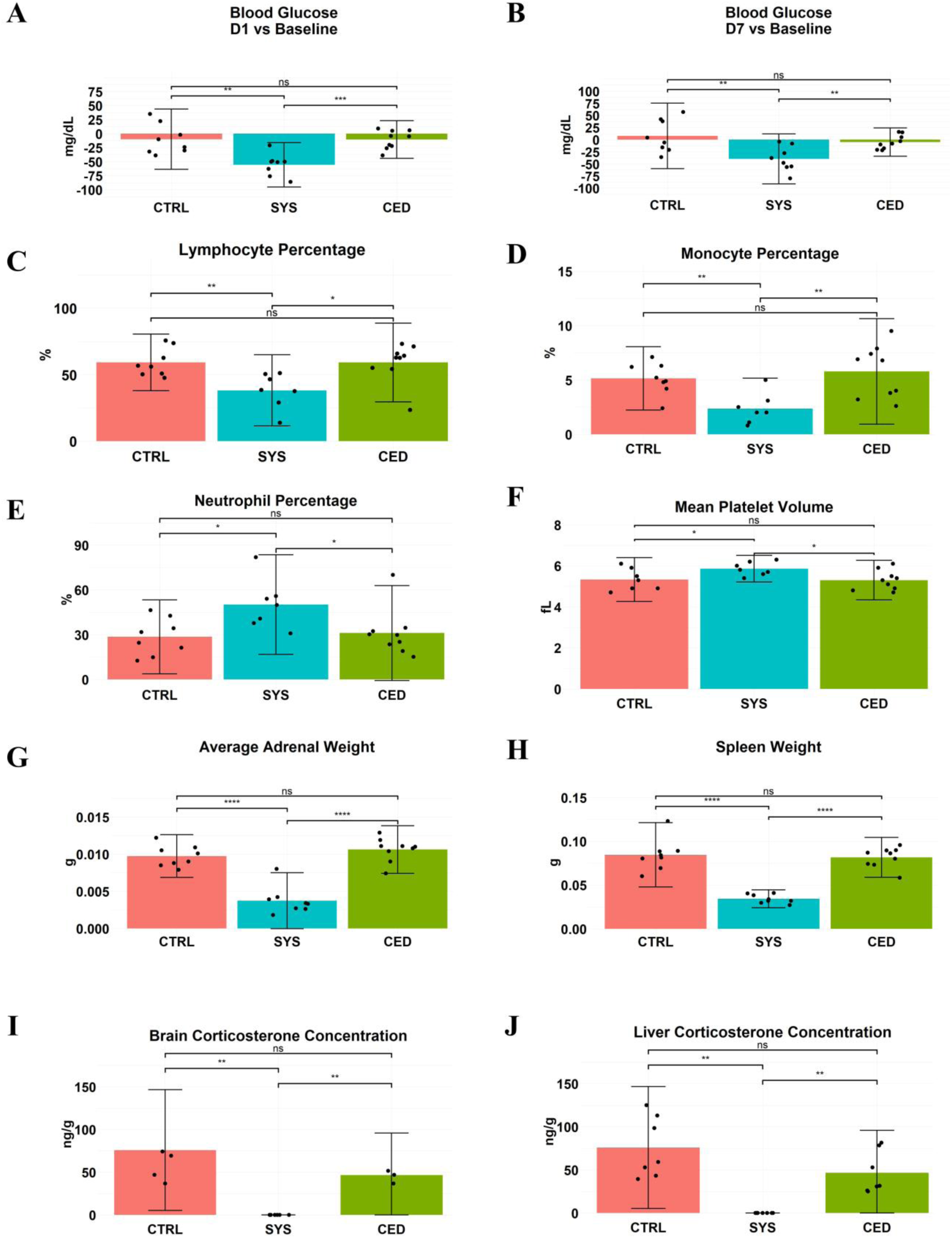
Systemic dexamethasone, but not CED, induces systemic side effects changes consistent with glucocorticoid toxicity. **A-B** – Systemic dexamethasone lowered blood glucose relative to pretreatment baseline at day 1 and day 7, whereas CED-dexamethasone did not. On day 1, systemic dexamethasone reduced glucose versus CTRL and CED (Kruskal-Wallis p = 0.002; post-hoc t-tests p = 0.0019; p = 0.00018, respectively), while CED did not differ from CTRL (p = 0.95). On day 7, systemic dexamethasone remained lower than CTRL and CED (Kruskal-Wallis p = 0.01; post-hoc t-tests p = 0.0075; 0.0066, respectively), whereas CED again did not differ from CTRL (p = 0.34). **C-E** – Systemic dexamethasone altered circulating leukocyte composition compared to control and CED mice, respectively, after 7 days. Lymphocytes were reduced (CTRL: 59 ± 21%, CED: 59 ± 30%, SYS: 38 ± 27%; Kruskal-Wallis p = 0.008; post-hoc t-tests p = 0.0062 and p = 0.01, respectively). Monocytes were reduced (CTRL: 5.1 ± 2.9%, CED: 5.8 ± 4.9%, SYS: 2.4 ± 2.4%; Kruskal-Wallis p = 0.01; post-hoc t-tests p = 0.0024 and p = 0.0035, respectively). Neutrophils were reduced (CTRL: 29 ± 25%, CED: 31 ± 31%, SYS: 50 ± 33%; Kruskal-Wallis p = 0.02; post-hoc t-tests p = 0.017 and p = 0.038, respectively). CED did not differ from CTRL (p = 1, p = 0.51, and p = 0.72, respectively). **F-H** – Systemic dexamethasone also changed mean platelet volume and reduced adrenal and spleen weights compared to control and CED mice, respectively, after 7 days. Mean platelet volume differed (CTRL: 5.3 ± 1.1 fL, CED: 5.3 ± 0.9 fL, SYS: 5.9 ± 0.6 fL; Kruskal-Wallis p = 0.06, post-hoc t-tests p=0.05 and p 0.016, respectively). Average adrenal weight decreased (CTRL: 0.010 ± 0.003 g, CED: 0.011 ± 0.003 g, SYS: 0.004 ± 0.004 g; Kruskal-Wallis p = 0.0004; post-hoc t-tests p < 0.0001 and p < 0.0001, respectively). Spleen weight decreased (CTRL: 0.084 ± 0.037 g, CED: 0.082 ± 0.022 g, SYS: 0.034 ± 0.010 g; Kruskal-Wallis p = 0.0004; post-hoc t-tests p < 0.0001 and p < 0.0001, respectively). CED did not differ from CTRL (p = 0.91, p = 0.25, and p = 0.70, respectively). **I-J** – Endogenous corticosterone concentrations were ablated by systemic dexamethasone but preserved with CED. Liver corticosterone was 75.9 ± 35.4 ng/g (CTRL), 46.6 ± 46.6 ng/g (CED), and SYS: 0 ± 0 ng/g (SYS); brain corticosterone concentrations were 56.8 ± 36.0 ng/g, 45.1 ± 15.2 ng/g, and 0 ± 0 ng/g, respectively. (* = p < 0.05; ** = p < 0.01, *** = p < 0.001, **** = p < 0.0001)

Systemically delivered dexamethasone also led to a significant decrease in lymphocyte percentages (CTRL: 59 ± 21%, CED: 59 ± 30%, SYS: 38 ± 27%) and monocyte percentages (CTRL: 5.1 ± 2.9%, CED: 5.8 ± 4.9%, SYS: 2.4 ± 2.4%) after seven days of treatment (Figure 3C, D). Similarly, neutrophil percentages (CTRL: 29 ± 25%, CED: 31 ± 31%, SYS: 50 ± 33%) and mean platelet volume (CTRL: 5.3 ± 1.1 fL, CED: 5.3 ± 0.9 fL, SYS: 5.9 ± 0.6 fL) were increased following systemic treatment (Figure 3C, D). No significant alterations in blood counts were observed when dexamethasone was given by CED (Figure 3C, D, E, F). Additionally, splenic and adrenal atrophy were observed after 7 days of systemic delivery of dexamethasone and not with CED (Figure 3G, H). In line with the above findings, we observed a complete ablation of endogenous corticosterone in the brain and liver in mice with systemic dexamethasone that was not observed with CED of dexamethasone (Figure 3I, J).

### Analysis of murine tissue immediately after CED of dexamethasone shows a marked anti-inflammatory effect

In vitro studies showed that the concentration of dexamethasone delivered by CED is below the level affecting murine tumor cell viability (Supplementary Figure 1A). Therefore, we performed post-treatment tissue analyses to determine the effects of CED of dexamethasone on the tumor microenvironment. Histological analysis and snRNA-sequencing of tissue were performed immediately following seven days of CED of dexamethasone or vehicle with an Alzet osmotic pump, starting at 21 days post injection. A systemic delivery group was included for histological analysis utilizing dosages used in previous studies of glioma-bearing mice (10 mg/kg/day i.p. dexamethasone) (8, 17, 23, 24) (Figure 4A).

**Figure 4:**
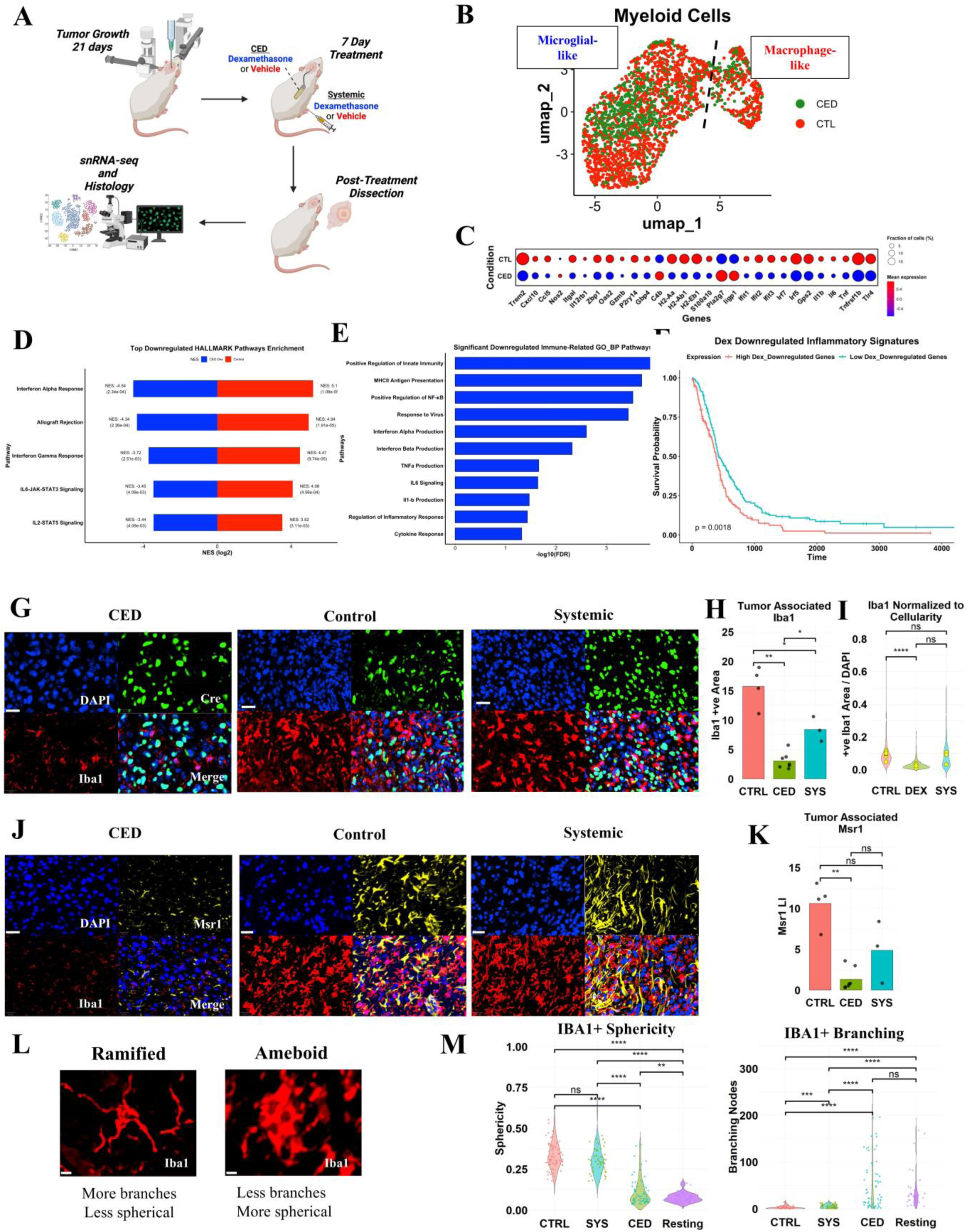
CED of dexamethasone suppresses the inflammatory myeloid program in glioma, reduces Iba1/MSR1-associated myeloid cells, and shifts tumor-associated myeloid cells toward a less activated morphology. **A** – Schematic of the post-treatment tissue analysis experiment. Mice were implanted with tumors on DPI 0. On DPI 21, mice received either CED of dexamethasone (100 ng/μl), daily systemic i.p. injections of dexamethasone (10 mg/kg), or PBS vehicle. All mice received both injections and pump implantation. After seven days of treatment, pumps were explanted, and the mice were sacrificed for either snRNA-seq or histology. **B** – UMAP of myeloid nuclei showing macrophage-like (Msr1+, Cd74+) and microglial-like (P2ry12+, Tmem119+) clusters. **C** – CED of dexamethasone reduced expression of a broad set of immune activation genes in myeloid nuclei. **D-E** – GSEA of Hallmark and GO biological pathways showed inflammatory and immune-related pathways were positively enriched in vehicle-treated mice and negatively enriched after CED-dexamethasone. Cytokine signaling pathways including IL6, IL-1β, and TNFα, were significantly downregulated. **F** – A survival analysis utilizing TCGA and CGGA bulk RNA-seq showed that greater-than-median expression of the top 50 significantly downregulated myeloid genes following CED-dexamethasone was correlated with worse overall survival (p = 0.0018). **G** – Representative immunofluorescence images showing Iba1 positivity in Cre-positive fields. Cre positivity is specific to murine tumor cells; scale bar = 20 μm. **H** – Iba1-positive staining area was decreased throughout tumors in dexamethasone-treated mice (CTRL vs SYS, p = 0.018; CTRL vs CED, p = 0.0031; CED vs SYS, p = 0.031). **I** – Iba1-positive staining area remained reduced with CED-dexamethasone after normalization to total cellularity as a proxy for tumor burden (p < 0.0001). **J** – Representative immunofluorescence images showing Iba1 and Msr1 in the tumor core; scale bar = 20 μm. **K** – CED-dexamethasone reduced Msr1-positive cells versus control (p = 0.0032); systemic dexamethasone showed a nonsignificant trend (p = 0.24). **L** – Representative ramified and ameboid myeloid cell; scale bar = 5 μm. **M** – Automated morphological analysis of tumor-associated, Iba1-positive myeloid cells across treatment groups and contralateral, “resting” brain showed CED-dexamethasone made tumor-associated myeloid cells less spherical and more branching; statistics computed per microglia. (* = p < 0.05; ** = p < 0.01, *** = p < 0.001, **** = p < 0.0001)

We performed snRNA-sequencing of tumor-containing brain tissue from the CED-dexamethasone-treated and vehicle-treated samples. These nuclei were projected in UMAP space and assigned cell lineages using canonical marker genes, which were validated with SingleR (Supplementary Figure 3A, B). Shifts in transcriptional patterns between treatment conditions in myeloid cells, the predominant inflammatory cell type captured in our analysis, were assessed. UMAP visualization demonstrated that myeloid cells clustered into resting microglial (*P2ry12*, *Hexb*, *Sall1*) and activated macrophage/monocyte-like subpopulations (*Msr1, Cd206/Mrc1, Lyz2*) (Figure 4B; Supplementary Figure 3C) (44). Microglial-like nuclei were enriched in *P2ry12* and *Hexb* expression, while the macrophage/monocyte-like nuclei were enriched in *Msr1* (Supplementary Figure 3C). Analysis of a broad set of inflammatory genes showed that CED of dexamethasone reduced reactive microglial and macrophage genes, including markers of interferon signaling (*Irf7, Ifit2, Ifit3*) (45), phagocytosis and antigen presentation (*H2-Aa, H2-Ab1*) (46), and chemokine/cytokine signaling (*Tlr4, Il1b, Il6, Tnf*) (47) (Figure 4C).

The anti-inflammatory transcriptional effects of CED of dexamethasone were further examined by performing gene set enrichment analysis (GSEA) (48) on our treatment groups. Analysis of significant Hallmark pathways revealed that the most significantly downregulated pathways were all related to inflammation, including interferon response, IL2-STAT5 and IL6 signaling, and immune rejection pathways (Figure 4D). GSEA analysis of Gene Ontology (GO) pathways provided further resolution, highlighting the suppression of several inflammatory and immune activation pathways. Pathways involving positive regulation of innate immunity, MHCII antigen processing, inflammatory cytokine production and signaling, interferon production and signaling were all downregulated with CED of dexamethasone (Figure 4E).

To assess the prognostic importance of the reduction of inflammatory transcriptional signatures, we performed log-rank testing on the human homologues of the top 50 significantly downregulated myeloid cell genes after CED of dexamethasone (Supplementary Table 2). Using VST normalized TCGA and CGGA bulk sequencing data from IDH-WT primary GBM patients (25), a significantly improved survival in patients with low expression of these inflammatory genes was observed (log-rank, p=0.0018; Figure 4F).

Immunohistochemical staining of post-treatment tumors demonstrated a significant decrease in Iba1 staining within the tumor of mice treated with CED dexamethasone compared control mice and mice treated with systemic dexamethasone (Figure 4G, H). This decrease in Iba1 positivity was also observed when normalized to tumor cellularity (Figure 4I). We also examined Msr1, a marker of tumor-associated macrophages that has been associated with a poor prognosis in glioblastoma (49, 50) (Figure 4J). We observed that Msr1 labeled a distinct subset of myeloid cells within the tumor core, which was non-overlapping with P2ry12-positive microglia at the tumor margin (Supplementary Figure 4). This supports the idea that Msr1 is labeling a subset of “activated” myeloid cells that emerge in the setting of chronic tumor-inflammation. We found a significant decrease in Msr1-positive cells with CED dexamethasone over control, whereas the effects of systemic dexamethasone were not significant (Figure 4K).

In addition to a decrease in the overall abundance of Iba1-positive and Msr1-positive myeloid cells after CED of dexamethasone, the morphology of Iba1-positive cells was significantly more ramified in mice treated with CED of dexamethasone. It has been suggested that microglial morphology (i.e., a hypertrophic, amoeboid appearance) is a sign of activated microglia, while resting microglia take on a ramified appearance (51–53). Using a previously developed and validated automated morphological analysis toolkit (54), we quantified sphericity and branching in a subset of Iba1-positive cells from mice treated with systemic dexamethasone, CED dexamethasone, and vehicle. Tumor-associated Iba1-positive cells were less spherical and had more branching nodes in mice treated with CED of dexamethasone than in the systemic- or vehicle-treated conditions (Figure 4L).

To explore whether vascular changes might contribute to changes in myeloid recruitment, we also quantified CD34-positive tissue across treatment groups (Supplementary Figure 5A). CD34-positive area normalized to cellularity was reduced in CED-treated tumors relative to controls (Supplementary Figure 5B). Systemic dexamethasone showed an intermediate effect, with an average CD34-positive area lower than controls, but higher than CED-treated tumors, although pairwise differences with systemic-dexamethasone were not statistically significant. A similar pattern was observed in end-stage tumors from our GL261 survival study (Supplementary Figure 5C). These findings suggest that prolonged local dexamethasone therapy leads to an overall reduction in tumor vascularity to a greater degree than systemic drug delivery. Post-treatment snRNA-seq supports this finding, showing a significant reduction in *Vegfa* expression in astrocytes following CED-dexamethasone. Systemic dexamethasone did not lead to a significant transcriptional suppression of *Vegfa* in astrocytes (Supplementary Figure 3D).

We investigated the differential effects of local versus systemic dexamethasone therapy by comparing snRNA-seq data from myeloid cells in each group. Differential gene expression revealed 463 differentially expressed genes (Supplementary Table 3), many of which are inflammatory in ontology. Genes such as Cd68 and Ccr2 are significantly downregulated in the CED group compared to systemic administration, consistent with the broad anti-inflammatory effect observed with local delivery. Immunofluorescence staining for Cd68 (Supplementary Figure 5D) supported this transcriptional observation. The Cd68 labeling index was markedly reduced in the CED group relative to both control and systemic dexamethasone groups, whereas systemic treatment showed only a partial reduction relative to control (Supplementary Figure 5E). This is consistent with the observed marked decrease in Iba1-positive myeloid cells seen earlier. Not all inflammatory genes are downregulated, however, as evidenced by an upregulation in Ly86 and Stat1 in CED-dexamethasone mice compared to systemic-dexamethasone (Supplementary Table 3). Prolonged high levels of dexamethasone achieved during local delivery may lead to different transcriptional changes than the peak-trough pattern achieved by systemic drug delivery.

Comparison of enrichment scores in Hallmark pathways between the systemic and local groups showed a consistent negative enrichment in inflammatory pathways in the CED group. Systemic delivery, on the other hand, showed a similar negative enrichment in some Hallmark pathways (e.g., interferon response and allograft rejection), whereas a positive enrichment was observed in some key inflammatory ontologies (e.g., IL6 and TNFα signaling) (Supplementary Figure 6A). Finally, we assessed the clinical significance of RNA expression patterns in our CED and systemic groups by comparing them to recently published myeloid cell expression programs in glioblastoma (55). We found that systemic and local dexamethasone caused opposite enrichments (positive and negative, respectively) in two of these expression programs – “complement immunosuppressive,” and “systemic inflammatory.” (Supplementary Figure 6B)

### In vitro treatment of iPSC-derived microglia with dexamethasone inhibits LPS-induced activation

To assess the direct effects of dexamethasone on microglia cells in vitro, we utilized human induced pluripotent stem cell (iPSC) derived microglia (Figure 5A) (56), which are thought to more closely represent resting, in vivo microglia than immortalized myeloid cell lines (57, 58). Bulk RNA sequencing was analyzed from iPSC-derived microglia treated for 24 hours with dexamethasone or lipopolysaccharide (LPS), either alone or in combination.

**Figure 5:**
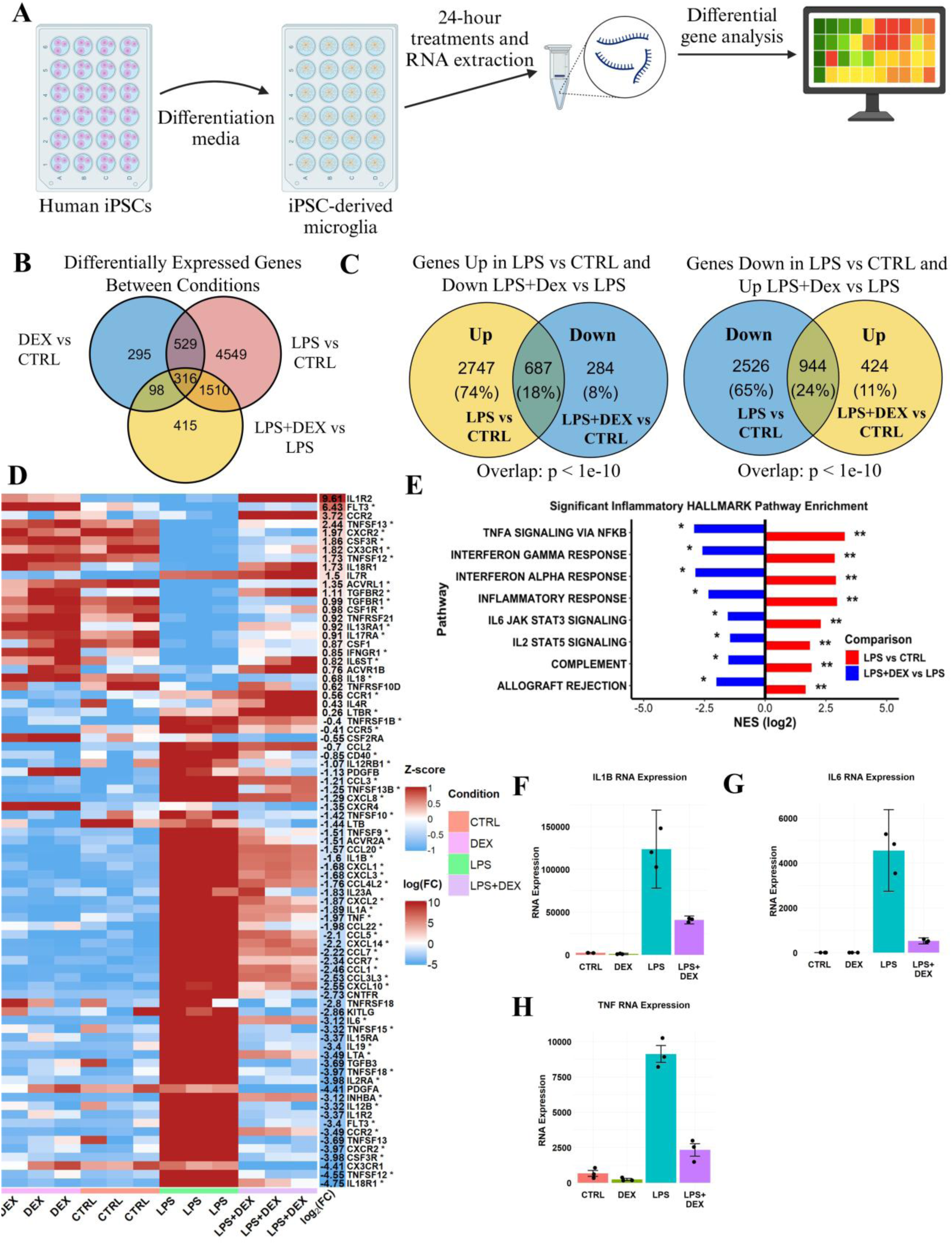
Dexamethasone broadly reverses the LPS-induced inflammatory transcriptional program in iPSC-derived microglia, including cytokine expression and Hallmark inflammatory pathway activation. **A** – Schematic of the iPSC-derived microglial experiment. iPSC cells were differentiated into microglia before being treated for 24 hours. RNA was then extracted from the cells differential gene expression analysis was performed. **B** – Venn diagram representing the number of unique and overlapping differentially expressed (p-adj < 0.05) genes in each comparison. **C** – The addition of dexamethasone to LPS reversed the expression of a large number of genes that were initially differentially expressed with LPS. The number of overlapping genes that were differentially expressed in opposite directions in the LPS vs CTRL and LPS+Dex vs LPS comparisons was significantly greater than would be expected by chance (p < 1e-10, hypergeometric test). **D** – Dexamethasone broadly reversed a large number of cytokines that were upregulated with the addition of LPS. The log_2_(FC) column represents the log-fold change in expression between the LPS and LPS+Dex conditions. An asterisk (*) next to a gene name denotes that LPS induced a significant change in expression that was significantly reversed in the LPS+Dex condition. **E** – LPS induced a broad upregulation of Hallmark inflammatory genes, which were relatively downregulated when cells were treated with a combination of LPS+Dex. **F, G, H** – Normalized RNA expression for IL-1β, IL6, and TNFα.

Differentially expressed genes between treatment conditions were calculated (Figure 5B). We found that LPS was a robust inflammatory stimulus with over 6,000 differentially expressed genes compared to control cells (padj < 0.05). A significant overlap was observed between genes differentially expressed in the LPS plus dexamethasone versus LPS comparison and genes differentially expressed in the LPS versus control condition. Namely, a significant number of genes that were significantly upregulated with LPS were significantly downregulated with the addition of dexamethasone and vice versa (Figure 5C). The pool of overlapping genes was significantly greater than would be expected by random chance (p < 1 ×10^−10^ by hypergeometric testing).

Similarly, LPS significantly altered the transcription of a broad range of cytokines. When dexamethasone was added to LPS, cytokine transcription typically shifted back toward control levels (Figure 5D, F-H). Additionally, analysis of Hallmark Pathways shows significant enrichment in inflammatory pathways (e.g., complement, TNFα, interferon, IL6, and IL2 signaling pathways) that are correspondingly negatively enriched with the addition of dexamethasone in combination with LPS (Figure 5E).

To confirm that dexamethasone-mediated suppression of inflammatory cytokines generalizes across human microglial and macrophage models, we performed qPCR in HMC3 microglia with vehicle, LPS, dexamethasone, or LPS+dexamethasone. LPS significantly increased RNA expression of IL1B, TNF, and induced a trend toward increased IL6 after 24 hours. Cotreatment with dexamethasone significantly attenuated IL1B, TNF, and IL6 RNA expression compared to LPS treatment alone, recapitulating what was observed in iPSC-derived microglia (Supplementary Figure 7A). Similar patterns were also observed in THP-1 cells using published bulk RNA sequencing data from Ansari et al (59). LPS induced expression of Hallmark inflammatory gene sets, which was attenuated by the addition of dexamethasone (Supplementary Figure 7B). LPS significantly increased RNA expression of IL1B, TNF, and induced a trend toward increased IL6 after 24 hours. Cotreatment with dexamethasone significantly attenuated IL1B, TNF, and IL6 RNA expression compared to LPS treatment alone, recapitulating what was observed in iPSC-derived microglia and HMC3 microglia (Supplementary Figure 7C). This suggests that the anti-inflammatory patterns of dexamethasone are consistent among a variety of myeloid cell subsets.

## Discussion

Dexamethasone is the main therapeutic for controlling glioma-associated edema, but its clinical efficacy is diminished by systemic toxicities. In this study, we used CED of dexamethasone to achieve high doses of dexamethasone in the brain with negligible systemic exposure. CED of dexamethasone led to enhanced intratumoral drug penetration and eliminated the metabolic and hematologic side effects seen with systemic drug delivery. Also, CED of dexamethasone suppressed tumor-associated inflammation to a greater extent than systemic delivery. While local dexamethasone therapy led to a modest survival benefit in our PDGFA-driven model, this effect did not generalize to GL261. Importantly, CED of dexamethasone was well tolerated across models.

CED of dexamethasone profoundly reduces inflammation in the tumor microenvironment. This anti-inflammatory effect involves broad transcriptional suppression of inflammatory pathways as well as a decrease in Iba1-positive myeloid cells in and around the tumor. Iba1-positive cells inside CED-treated tumors were also significantly more ramified in their appearance. Ramified microglia have often been interpreted as more resting, while a morphological shift towards an amoeboid appearance is a sign of their response to an inflammatory stimulus (51, 60, 61). This suggests that dexamethasone has direct effects on microglial activation. Our iPSC-derived microglial data validated the direct effects of dexamethasone on microglia, demonstrating that LPS-induced activation was partially suppressed by concurrent treatment with dexamethasone. We also found a significant depletion of Msr1-positive macrophages along with a reduced Cd68 labeling index in tumors treated with CED-dexamethasone, supporting the idea that local delivery of dexamethasone leads to a broad suppression of glioma-associated inflammation in both microglia and macrophages. Importantly, this reduction in inflammation was greater in magnitude and more consistent than what was observed with systemic dexamethasone delivery. We also found that lower expression of the top inflammatory myeloid genes downregulated by CED was associated with improved survival in the TCGA and CGGA GBM cohorts. Although correlative, this suggests that the inflammatory transcriptional program suppressed by local dexamethasone is clinically relevant and associated with adverse clinical outcomes.

Additionally, comparative snRNA-seq analysis between the CED and systemic murine groups revealed unique differences in inflammatory gene expression depending on the route of drug administration. Recently, Miller et al. noted that a unique inflammatory gene expression pattern, termed “complement immunosuppressive,” was upregulated in human patients receiving systemic dexamethasone therapy. In line with this, we found a positive enrichment of this pathway in mice treated with systemic dexamethasone and negative enrichment of the pathway in mice treated with CED dexamethasone. More broadly, local and systemic dexamethasone produced distinct inflammatory transcriptional states in myeloid cells, with greater suppression of genes such as Cd68, Msr1, and Ccr2, and more consistent negative enrichment of inflammatory ontologies. These results suggest that local dexamethasone both inhibits local microglial cell activation and reduces Msr1+ macrophages in the tumor core in a way that depends on the route of drug administration. Future work will have to elucidate the specific mechanisms of local dexamethasone’s anti-inflammatory properties in the setting of glioma.

CED of dexamethasone effectively avoided systemic toxicities while inhibiting tumor-associated inflammation. Mice treated with CED of dexamethasone exhibited stable blood glucose levels, no changes in hematologic measurements, and no evidence of splenic or adrenal atrophy, all of which were observed in mice receiving systemic dexamethasone therapy. The decrease in monocyte and lymphocyte percentages in the blood with systemic dexamethasone mirrors reported human side effects, as does the increase in neutrophil percentage and mean platelet volume (62, 63). We also observed an acute decrease in blood glucose following i.p. dexamethasone administration, which has been observed in mice by other groups, although its mechanism remains unclear (64, 65). Additionally, systemic dexamethasone eliminated endogenous corticosterone production in our mice, while corticosterone levels were unaffected with local delivery of dexamethasone.

Systemic dexamethasone administration led to high concentrations of the drug in the liver with relatively poor partitioning in the brain parenchyma. This aligns with other murine studies that have shown that dexamethasone is a substrate for the p-glycoprotein efflux pump encoded by Mdr1a on the blood-brain barrier and is actively transported out of the brain parenchyma (66, 67).

CED of dexamethasone delivers the drug directly into the brain parenchyma, which allows for better partitioning of the drug in the brain compared to peripheral organs. Through continuous infusion, CED leads to a roughly constant concentration of drug throughout the treatment period (33, 68), compared to peaks and troughs with systemic dosing (69). Dexamethasone had a short half-life in our albino B6 mice, approximately 1.1 hours, which is faster than the half-life of 2.3 hours reported in Wistar rats and 4 hours in human plasma (70, 71). We also showed that CED of dexamethasone leads to a significantly higher 24-hour AUC, making it far more effective than systemic delivery. Dexamethasone’s relatively rapid clearance from the brain parenchyma is likely responsible for the far more potent anti-inflammatory effects seen with CED, which constantly infuses new drug into the parenchyma, over systemic delivery. Continuous local exposure may sustain anti-inflammatory signaling more effectively than intermittent systemic peak-trough dosing.

A prior study by Nestler *et al.* assessed the degree to which systemically delivered dexamethasone accumulates in the tumor microenvironment in human glioblastoma patients (72). Nestler *et al.* report a mean brain tumor dexamethasone concentration of 225 ng/g, nearly ten-fold higher than we found in our contrast-enhancing biopsies. Importantly, Nestler *et al.* included tumor samples from patients with gliomas, meningiomas, and metastases who received 12-24 mg/day of dexamethasone for several days prior to their operation. In contrast, we collected biopsies from glioblastoma patients who received a 10 mg dexamethasone bolus as part of standard of care for a craniotomy, to examine how dexamethasone partitions between tumor tissue and plasma. Drug partitioned more effectively into contrast-enhancing tumor biopsies compared to non-enhancing biopsies, suggesting that an intact blood-brain barrier limits dexamethasone’s brain penetrance. Thus, the non-enhancing tumor that remains post-resection achieves the lowest concentration of systemic dexamethasone, providing additional justification for CED of the drug.

In addition to suppressing inflammatory myeloid programs, CED of dexamethasone also appeared to reduce tumor vascularity. We observed decreased CD34-positive area in CED-treated tumors, along with reduced *Vegfa* transcription in astrocytes, whereas systemic dexamethasone was ineffective in reducing *Vegfa* transcription and had a statistically insignificant intermediate effect on tumor vascularity. Because *Vegfa* can influence edema, vessel permeability, and myeloid recruitment (73), the observed reduction in tumor vascularity may represent an additional mechanism by which CED suppresses tumor-associated inflammation. Dexamethasone is a pleiotropic drug, and its effects on vascular and myeloid cells highlight that its mechanism is shaped through complex interactions among a variety of cell types.

The findings in this study have potential direct translational relevance for patients with glioblastoma, particularly those who remain dependent on dexamethasone for prolonged control of tumor- or treatment-related edema. Currently, many patients require extended systemic dexamethasone therapy, which is limited by cumulative toxicities. Our results suggest that CED of dexamethasone could provide a route to administering steroid therapy that spares systemic tissues. This strategy is particularly plausible given recent advances in chronic CED using implantable, refillable pumps, which permit prolonged and repeated intracerebral infusions without repeat surgeries (41). In this context, CED-dexamethasone could potentially be given to patients with recurrent GBM requiring dexamethasone or integrated with other CED-based therapies. Future clinical studies will be needed to define the optimal dose, infusion, schedule, and safety of long-term local dexamethasone delivery, as well as its effects on symptom control, quality of life, and compatibility with concurrent anti-tumor therapies.

## Methods

### Cell Viability Assays

For murine tumor cell lines (APCL and GL261), viability after dexamethasone treatment was assessed using the CellTiter 96® AQueous One Solution MTS Assay (Promega, G3582). For human tumor cell lines, viability was assessed using the CellTiter-Glo® Luminescent Cell Viability Assay (Promega, G7570). All cell lines are confirmed to be mycoplasma free using ATCC Kit 30-1012K. In both cases, cells were plated in the inner wells of 96-well plates, allowed to settle overnight, and then treated with the indicated concentrations of dexamethasone for 72 hours. All conditions were performed in triplicate. Assays were developed according to the manufacturer’s instructions, and absorbance at 490 nm (MTS) or luminescence (CellTiter-Glo) was measured using a plate reader. IC50 values were calculated using the drc package in R.

### Animal Ethics Statement

All experiments were performed in accordance with institutional and national guidelines for the care and use of laboratory animals. Experimental protocols were approved by the Institutional Animal Care and Use Committee (IACUC) at Columbia University. Female B6(Cg)-Tyr^c-2J^/J mice from Jackson Laboratories (strain #000058, age 6-8 weeks) were used for all experiments. Mice were housed under standard, pathogen-free conditions with a 12-hour light/dark cycle and had unrestricted access to food and water.

### Orthotopic Cell Injections

Our p53^-^/^-^ PDGFA+ murine glioma cell line (APCL) was generated as previously described (74). GL261-Luc was previously described in Kim et al., 2017 (75). Resulting cell lines were grown at 37°C with 5% CO2.

Orthotopic tumor implantation was performed as previously described (74). Briefly, mice were anesthetized with Ketamine/Xylazine (100 mg/kg and 10 mg/kg, respectively) and assessed for lack of reflexes by toe pinch. Hair was shaved and scalp skin incised. The skull was cleaned with a cotton swab and the bregma was identified. A burr hole was made with a 17-gauge needle 2 mm lateral and 2 mm anterior to the bregma. A cell suspension was made from lifted cell lines. An intracranial injection was performed under stereotactic guidance, 2 mm deep into the brain parenchyma using a Hamilton syringe at a flow rate of 0.3 μl/min to deliver 50,000 APCL cells or 6,000 GL261 cells with a volume less than 2 μl. Tumor growth was assessed through bioluminescence imaging as previously described (76). Mice were approximately 8 weeks old when they were injected with tumors.

### CED of Dexamethasone and Systemic Administration

Pump implantation was performed as previously described (76). Alzet osmotic pumps (1007D) were filled with either PBS (control) or PBS with 100 ng/μl of dexamethasone-phosphate. Pumps were implanted at 21 DPI for all studies, and the catheter was implanted along the same burr hole that was used for tumor implantation. For mice receiving systemic administration of dexamethasone, a PBS-containing pump was implanted and mice received an i.p. injection of 10 mg/kg of dexamethasone daily. One percent Omniscan was included in all Alzet pumps to confirm successful drug distribution by MRI.

### Murine MRI Scans

MRI scans were conducted using a Bruker BioSpec 9.4T scanner (Bruker Corp., Billerica, MA). Mice were anesthetized with 1–2% isoflurane mixed with medical air and administered via a nose cone. The isoflurane concentration was adjusted throughout the scan to maintain a stable respiratory rate between 40 and 70 breaths per minute. Respiration was monitored using a sensor pillow connected to a physiological monitoring system (SA Instruments, Stony Brook, NY). To ensure consistent body temperature, a circulating water heating pad was used to maintain the temperature at approximately 37°C.

For initial localization, low-resolution T1-weighted scout images were acquired. High-resolution anatomical imaging was performed using a T2-weighted rapid acquisition with relaxation enhancement (RARE) sequence with the following acquisition parameters: repetition time (TR) = 3000 ms, echo time (TE) = 45 ms, field of view (FOV) = 17 × 15 mm, matrix resolution = 225 × 198, and slice thickness = 0.7 mm, with 16 slices spanning the entire brain. Contrast-enhanced imaging was performed using a T1-weighted sequence with the same geometric parameters but different acquisition settings: TR = 150 ms, TE = 2.2 ms, and flip angle = 70°.

### Murine Survival Studies

APCL survival study – Two survival studies were performed in the APCL model, each with 10 mice per group, and the survival results shown represent the combination of these two experiments. Tumors (50,000 cells) were implanted on DPI 0 and treatment began on DPI 21. Tumor growth was assessed by luciferase imaging. Mice that died during anesthetic overdose or that lacked luciferase signal were excluded from further analysis. Treatment with vehicle (PBS) or dexamethasone (100 ng/μl) lasted 7-days and pumps were removed on DPI 28. An MRI was conducted at this point to visualize intraparenchymal gadolinium from all pumps. Mice were followed until death or until tumors reached end stage as evidenced by peri-orbital hemorrhage, epistaxis, seizures, or severe lethargy.

GL261 survival study – One survival study was also performed in the GL261 model with control, systemic dexamethasone, and CED-dexamethasone groups, each with 10 mice. GL261 tumors (6,000 cells) were implanted on DPI 0, and treatment began on DPI 11. Tumor growth was assessed by luciferase imaging. One mouse died prior to pump implantation and was excluded from further analysis. Treatment with vehicle (PBS), daily i.p. dexamethasone 10 mg/kg or CED-dexamethasone (100 ng/μl via an Alzet 1007D pump) lasted 7-days, and pumps were removed on DPI 18. Injections and Alzet pump implantation were controlled for in all groups. An MRI was conducted at the time of pump removal to visualize intraparenchymal gadolinium from all pumps. Mice were followed until death or until tumors reached end stage as evidenced by peri-orbital hemorrhage, epistaxis, seizures, or severe lethargy.

### Post-treatment Tissue, and Dexamethasone Quantification Studies

For post-treatment analysis of tissue, mice were implanted with Alzet pumps on DPI 21, as described above. Following 7-days of treatment, pumps were removed, and mice were anesthetized and perfused with 15 mL PBS followed by 15 mL 4% PFA at a rate of 3 mL/min. Brains were excised and further fixed in 4% PFA for 24 hours. Brains were then embedded in paraffin and five μm thick sections obtained. Sections were stained following basic immunohistochemical methods. Following deparaffinization with xylene and ethanol, antigen retrieval was performed under pressure in a 10 mM, pH 6 citrate buffer with 0.05% tween-20. Sections were incubated in primary antibody solutions overnight at room temperature. Immunoperoxidase staining was accomplished with the appropriate Vector kit (PK-6100). Fluorescent secondary antibodies were used at 1:1000.

The following primary antibodies were used: Cre (1:50, Cell Signaling 15036), Iba1 (1:1000, Cell Signaling 17198), Iba1 (1:500, Aves 1BA1-100), MSR1 (1:500, Invitrogen PA5-102519), P2ry12 (1:500, BioLegend 848001), CD68 (1: 200, Abcam ab303565), and CD34 (1:200, Abcam ab81289). For standard immunofluorescence, fluorescent secondary antibodies included goat anti-rabbit Alexa Fluor 488 or 568 (A11008/A11036), goat anti-rat IgG2b Novus Biologics Fluor 488 (NB7129), and goat anti-chicken Alexa Fluor 647 (A21449). For CD68/CD34 staining, CD68 was applied first and detected using the Alexa Fluor™ 488 Tyramide SuperBoost™ Kit, goat anti-rabbit IgG (Invitrogen, B40922), followed by CD34 staining.

Images of immunoperoxidase staining were scanned at 40x using a Leica SCN 400 digital slide scanner. Immunofluorescence images were obtained on a Nikon AX confocal microscope. Images were quantified using QuPath (77).

For quantification of dexamethasone, dissected tissue or plasma was flash frozen in liquid nitrogen for later analysis (see *Quantification of Dexamethasone*). In mice receiving systemic dexamethasone, the mice were perfused with 20 mL of PBS before the tissue was dissected and frozen.

### Measurement of Physiologic Side Effects

Blood glucose measurements were taken by briefly sedating mice with isoflurane before pricking their tail with a 20-gauge needle. Glucose levels were determined with a Accu-Chek home diabetes glucose meter from Roche (78). Glucose levels were measured after 1-day, 2-days, and 7-days of treatment.

For murine CBCs, blood from the three treatment groups was collected via cardiac puncture immediately after CED pump removal prior to sacrifice and was placed into 0.5mL EDTA-anticoagulated tubes for CBC analysis. Samples (15uL) were analyzed using Heska Element HT5 Veterinary Hematology Analyzer (laser flow cytometry, colorimetric detection, and impedance technology). Maintenance and quality control procedures are performed daily. Samples were tested within one hour of collection. Samples submitted overnight and for repeated analysis (if initial counts were unanalyzable) were stored at 4°C and tested within 12 hours after acquiring. Before testing, blood samples were placed on a sample rocker and then gently inverted 4-5 times to ensure proper mixing. At this time, the samples were inspected for blood clots. Any samples with visible blood clots were rejected for analysis. Serum biochemistry was performed using the Heska Element DC5X Veterinary Chemistry Analyzer to measure glucose, triglyceride, total protein, creatinine, alanine aminotransferase (ALT), and aspartate aminotransferase (AST) levels. Fresh blood was collected from the same three groups and placed into a 0.8mL serum separator tube. Tubes were centrifuged at 2000g spin for 10 minutes after 15 minutes incubation at room temperature in the dark. The supernatant in the form of serum was transferred to a 0.5mL polypropylene tube specific to this analyzer. The samples were collected over 1 day, stored at - 20°C, and tested over two consecutive days. The serum was thawed at room temperature for 20-25 minutes before testing.

Adrenal and splenic atrophy was assessed by carefully dissecting the mouse spleen and adrenal glands from surrounding organs and adipose tissue. The mean weight of the adrenal glands was recorded along with the weight of the spleen. Corticosterone concentrations in homogenized liver and brain were measured by high-performance liquid chromatography-tandem mass spectrometry (HPLC-MS/MS).

### iPSC-derived Microglia Differentiation and Treatment

The differentiation of induced-pluripotent stem cells (iPSCs) to primitive hematopoietic progenitor cells (HPCs) is optimized using the STEMdiff™ Hematopoietic Kit (Catalog # 05310, STEMCELL Technologies) as adapted from McQuade et al.(58) iPSC microglial differentiation from HPCs was done through the following process. Cells were incubated at 37°C. Briefly, HPCs were plated on 24-well Cultrex coated plates at a density of 100,000 cells per well. Cells were cultured in DMEM/F12 (brand) medium, Insulin (ITS) 100x, B27 50x, N2 100x, Glutamax 100x, MEAA 100x, Monothioglycerol, Insulin 10mg/mL, PenStrep 100x. Cells were fed every 2 days with a tri-cytokine cocktail of these differentiation factors: 100ng/mL IL-34, 50ng/mL TGB1, and 25ng/ml M-CSF. On day 13, cells were gently lifted with mechanical disruption to split ∼50% of cells into new wells as confluency of cells allowed for distribution of cells into new plate for higher yield. Differentiation with the tri-cytokine mix continued every 2 days until day 25. On day 25, the addition of 100ng/mL CD200 and 100ng/mL CX3CL1 for further maturation and ensure homeostatic state. Feeding occurred daily until day 28 when the cells were ready for functional and transcriptomic assays. On day 28, the fully matured human iPSC microglia were treated with 25 ng/μl of dexamethasone (63 μM) or 1 ng/μl of LPS, either separately or in combination. The cells underwent RNA extraction as per the Qiagen reagent protocol as well as the creation of cDNA. Bulk RNA sequencing was then performed.

### HMC3 Microglia Treatment and Quantitative PCR; THP1 RNA Sequencing Analysis

Human HMC3 microglia were cultured under standard conditions in DMEM with 10% fetal bovine serum. Cells were plated in 6-well plates at a density of 3 x 10^5 cells per well. And treated for 72 hours with vehicle, LPS (1 ng/μl), dexamethasone (25 ng/μl), or the combination of LPS and dexamethasone. Each condition was performed in 3-4 biological replicates. Following treatment, RNA was extracted using the RNeasy Mini Kit (Qiagen, 74106). cDNA was synthesized using the Invitrogen SuperScript VILO cDNA Synthesis Kit according to the manufacturer’s protocol. Quantitative PCR was performed on a QuantStudio 6 Pro Real-Time PCR System using Thermo Scientific ABsolute Blue qPCR Mix, SYBR Green, Low ROX (AB4163A), with ACTB as the reference gene. The following primers were used: IL1B forward, ATGATGGCTTATTACAGTGGCAA, reverse, GTCGGAGATTCGTAGCTGGA; TNFA forward, CCTCTCTCTAATCAGCCCTCTG, reverse, GAGGACCTGGGAGTAGATGAG; IL6 forward, ACTCACCTCTTCAGAACGAATTG, reverse, CCATCTTTGGAAGGTTCAGGTTG; and ACTB forward, TGGCACCCAGCACAATGAA, reverse, CTAAGTCATAGTCCGCCTAGAAGCA. IL1B, IL6, and TNF expression was quantified by the ΔΔCt method, with fold change calculated as 2^-ΔΔCt relative to the CTRL group mean. Statistical analyses were performed on ΔCt values.

Bulk RNA sequencing data was obtained from (59) (GEO GSE208041) and analyzed using DESeq2(79). THP-1 cells were treated with vehicle control, LPS (100 ng/ml), and LPS+Dexamethasone (100 ng/ml + 1 μM) for 3 hours before RNA was extracted with TRIzol and sequenced on an Illumina NovaSeq 6000.

### Quantification of Dexamethasone

Dexamethasone was measured by high-performance liquid chromatography-tandem mass spectrometry (HPLC-MS/MS) on a platform comprising of Waters Xevo TQS triple quadrupole mass spectrometer integrated with a Waters Acquity UHPLC (Waters, Milford, MA). Tissue samples were homogenized in LCMS grade water using Precyllis lysing kits with a bead homogenizer (Benchmark, NH). Dexamethasone was extracted from the tissue homogenates and serum/plasma samples spiked with deuterated dexamethasone (d4 dexamethasone) by liquid-liquid extraction using tert-butyl methy ether (MTBE). Chromatographic separation was done on a Waters Aquity UPLC BEH C18 column (1.7µm, 2.1X50mm) maintained at 50°C employing gradient elution using water and methanol with 0.1% formic acid as mobile phases. Positive ESI-MS/MS mass spectrometry under MRM mode was performed using the following transitions: 393.30>355.21. Dexamethasone and 397.30>377.20 (d4 dexamethasone). The dexamethasone concentration was measured by comparing integrated peak areas against known amounts of dexamethasone using Targetlynx V4.2.

The limit of quantification of the assay is 0.5ng/ml. The mean intra-and inter-assay imprecision was 1.68% and 3.42% respectively.

### snRNA Sequencing of Murine Tissue

As above, mice were sacrificed following seven days of CED of dexamethasone or vehicle treatment. Mouse brains were quickly dissected and the tumor bearing quadrant of the brain was isolated. The quadrant was then flash frozen in liquid nitrogen for later analysis. Later, samples were homogenized and nuclei isolated according to the Chromium Single Cell Nuclei Isolation Kit protocol (1000494). Barcoding, library preparation, and sequencing were performed by the JP Sulzberger Columbia Genome Center. The fastq files were loaded into 10xGenomics cloud and mapped with Cell Ranger Count v9.0.0 onto Mouse library GRm39 2024-A.

Standard Seurat pipelines were used for the analysis of snRNA sequencing data (80). A feature/cell matrix (filtered) was used to create Seurat objects. The merged object was subset to include only those nuclei containing between 500 and 5000 features and mitochondria counts less than 5% across all samples. Doublets were called and excluded by using https://github.com/chris-mcginnis-ucsf/DoubletFinder. Normalization, scaling and integration of the data was performed with the Seurat SCTransform() function. Nuclei were projected into UMAP space, clustered, and assigned cell lineages. Clustering of nuclei was done using the shared nearest neighbor smart local moving algorithm, dimensionality PCA reduction using Seurat’s FindNeighbors() function. Major cell types were identified using canonical cell type markers and SingleR (25, 81). Gene set enrichment analysis (GSEA) was performed using standard downstream analysis methods for gene set enrichment workflows, including the R packages hypeR (“hypergeometric testing”) and the SingleSeqgSet package for a Wilcoxon Mann Whitney Correlation Corrected GSEA analysis. Gene sets included those from MSigDB C2 (KEGG, REACTOME, etc) and C5 (Gene Ontology) libraries. Only significant gene sets with an adjusted p-value < 0.05 were considered.

### Bulk RNAseq Survival Analysis and RNAseq Data Analysis

The count matrix for the TCGA GBM dataset was downloaded using the GDCquery tool in R. The Chinese Glioma Genome Atlas (CGGA) RNAseq datasets was downloaded from (http://www.cgga.org.cn/download.jsp) (82, 83). Counts were normalized using DESeq2 in R (79). Only primary IDH wildtype GBM samples were kept for downstream analyses (TCGA: 139 samples, CGGA: 179 samples). Survival analysis was performed using the survival package in R, using the enrichment of the CED-dexamethasone gene set. Kaplan-Meier survival analysis was completed using median expression values (high vs. low).

Bulk RNA sequencing data was normalized and differentially expressed genes between conditions were determined using DESeq2. Gene ontology of differentially expressed genes between iPSC conditions were determined using GSEA and Hallmark pathways.

### Statistics

Sample sizes were not predetermined by power analysis but were consistent with previous experiments from our laboratory. A minimum of n=3 biological replicates were utilized for all experimental conditions. Statistical analyses were performed in R (version 4.5.2). For comparisons involving more than two experimental groups, a Kruskal-Wallis H test was employed to assess global significance. When a significant difference was detected, pairwise comparisons were made using post-hoc Welch’s t-tests. For comparisons involving only two groups, a two-tailed Welch’s t-test was used. Statistics were computed on a per animal basis, except for IBA1-positive cell morphology which was computed on a per cell basis. Values are reported as mean ± standard error unless otherwise specified. A p-value of <0.05 was considered statistically significant.

### Sex as a biological variable

Our study exclusively examined female mice. It is unknown whether the findings are relevant for male mice.

## Data Availability

All murine snRNA sequencing and human iPSC bulk RNA sequencing datasets generated in this study were deposited to GSE326006. Values for all data points in graphs are reported in the Supporting Data Values file. All relevant materials are available with reasonable request to the corresponding author.

## Author Contributions

NWR, NBD, LL, and AJT are co-first authors. Experiments were designed by NWR, NBD, LL, AJT, PC, and JNB. NWR, NBD, LL, AJT, DET, NI, and MA carried out experiments. Survival studies and snRNA analysis were done by NBD LL, and AJT. Histological analysis was done by NWR, LL, and AJT. Mass spectroscopy experiments were done by NWR and LL. Analysis of bulk RNA data was done by NWR and AJT. NWR, NBD, and AJT wrote the initial draft of the manuscript which was reviewed by all authors. The final manuscript was approved by all authors. PC and JNB acquired funding and provided oversight. Co-first authors contributed equally to data generation and analysis. Final authorship order was the decision of the senior authors.

## Acknowledgements

We thank Dr. Renu Nandakumar and Jiyoung Kim from the Biomarkers Core Laboratory at the Columbia Irving Institute for Clinical and Translational Research, which is supported by the National Center for Advancing Translational Sciences, National Institutes of Health, through Grant Number UL1TR001873, for conducting the UPLC-MS/MS assays. We thank the staff at the Molecular Pathology Shared Research Core at Columbia University Irving Comprehensive Cancer Center for help embedding and sectioning tissue. These studies used the Confocal and Specialized Microscopy Shared Resource of the Herbert Irving Comprehensive Cancer Center at Columbia University, funded in part through the NIH/NCI Cancer Center Support Grant P30CA013696. The mouse MRI studies presented in this work were carried out in the MR Facility of the Oncology Precision Therapeutics and Imaging Core (OPTIC) Shared Resource which is supported by funds from the Columbia University Medical Cancer Center Support Grant (CCSG) and NHI grant #P30 CA013696 (National Cancer Institute).

