## Supplemental for "Convection-enhanced delivery of dexamethasone in glioma suppresses myeloid inflammation while avoiding systemic toxicities"

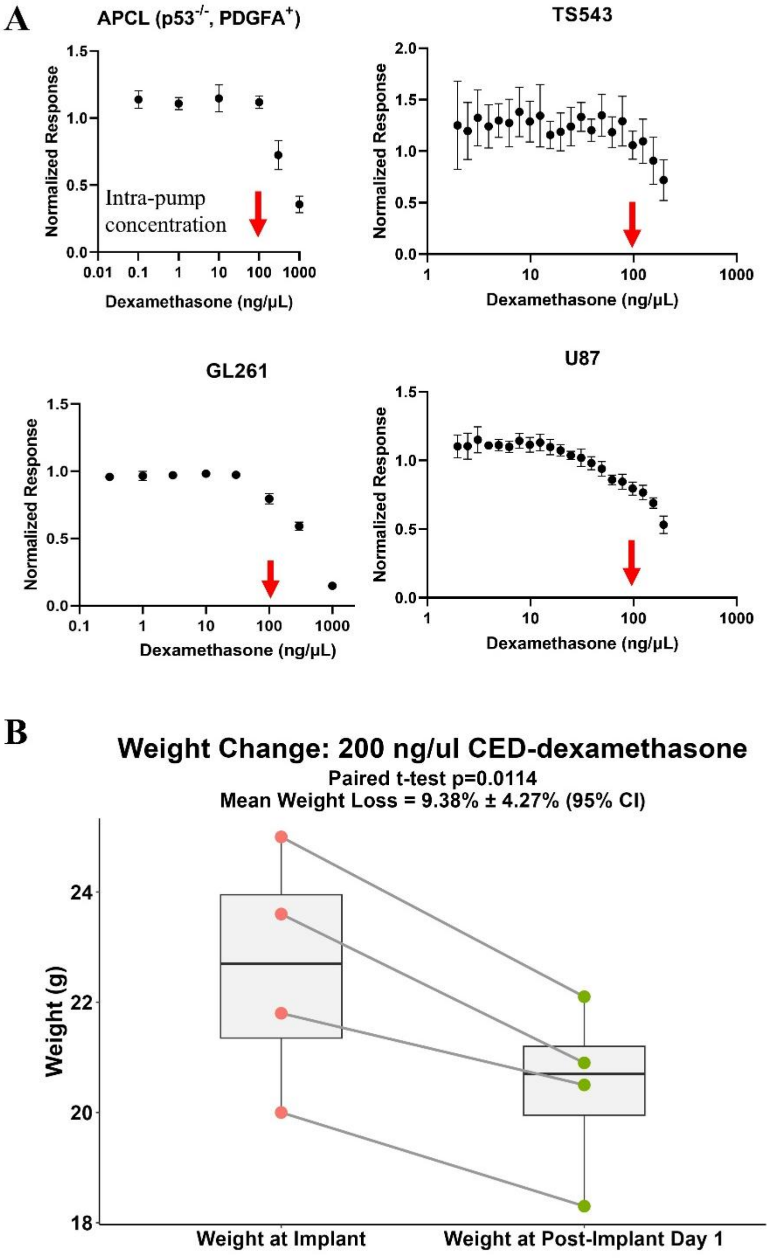


**Supplementary Figure 1**: **A** – The IC50 of dexamethasone was determined to be 549 ± 289 ng/µl (1,399 ± 736 µM) for murine APCL ($p53^{-/-}$, $PDGFA^{+}$) cells, 705 ± 219 ng/µl (1,796 ± 558 µM) in murine GL261 cells, 202 ± 19 ng/µl (515 ± 48 µM) in human TS543 glioma cells, and 194 ± 23 ng/µl (494 ± 59 µM) in human U87 cells. IC50 ± SE. **B** – Mice receiving CED-dexamethasone at a concentration of 200 ng/µl lost a significant amount of weight 1 day after pump implant (9.4% ± 4.27% (mean body weight loss ± 95% CI); paired t-test p = 0.01).

**A**

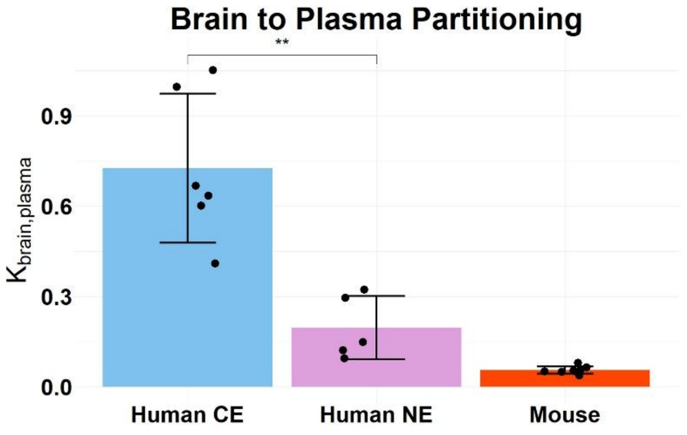


**Supplementary Figure 2**: **A** – The partitioning coefficient between brain and plasma in human contrast-enhancing tumor samples, non-enhancing samples, and mouse tumor-bearing brain quadrants. Contrast-enhancing tumor samples had brain to plasma concentration ratio of 73 ± 20% (mean ± 2*SE), while non-enhancing tumor samples had a brain to plasma concentration ratio of 20 ± 10% (Welch t-test, $p = 0.0021$). In our murine studies, the brain to plasma concentration in mice receiving systemic dexamethasone was 6 ± 2%.

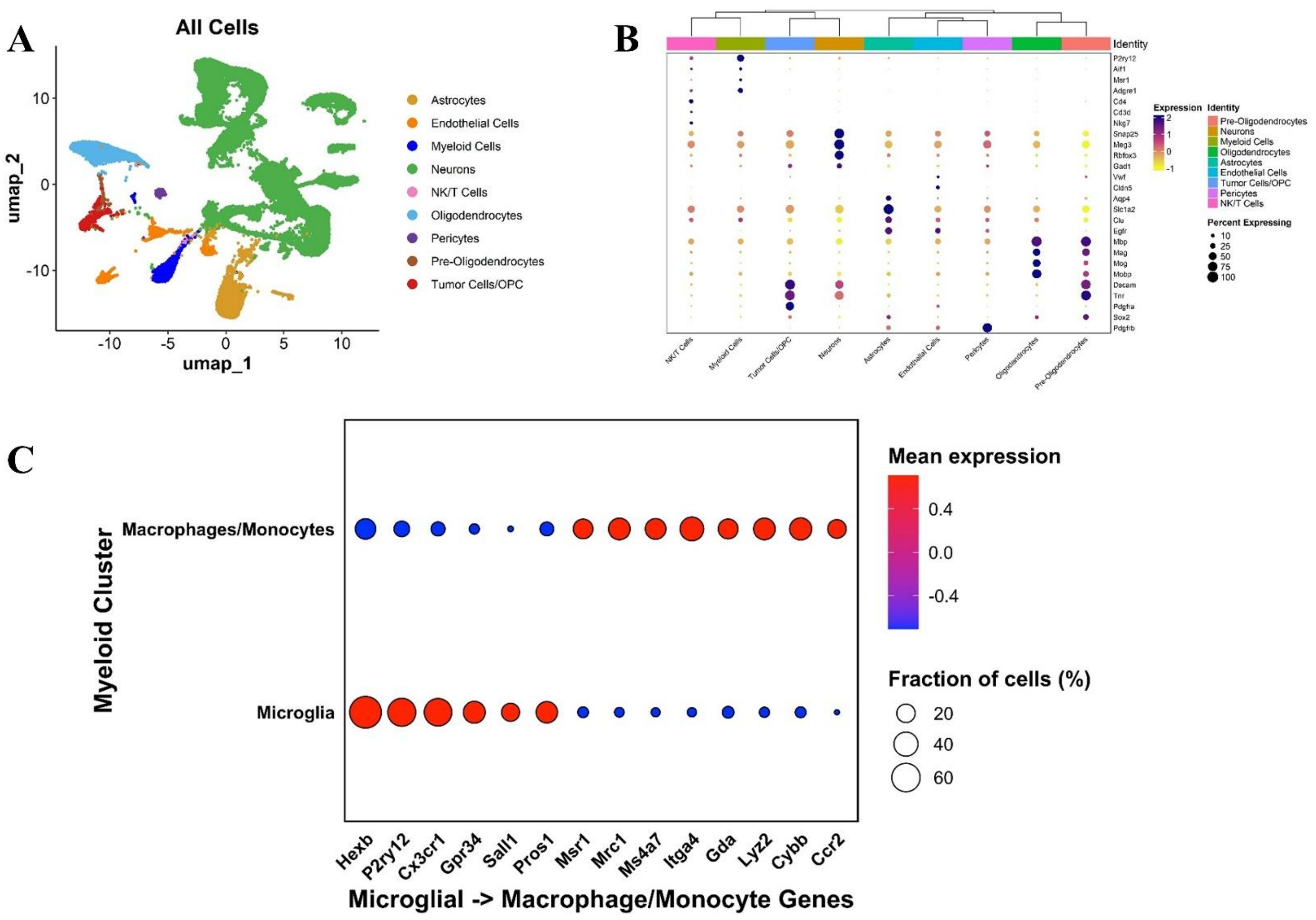


**Supplementary Figure 3**: **A** – Major cell types assigned by SingleR. **B** – Dot plot of canonical marker genes present in major cell lineages. **C** – Microglial and macrophage gene expression across myeloid nuclei using canonical genes.

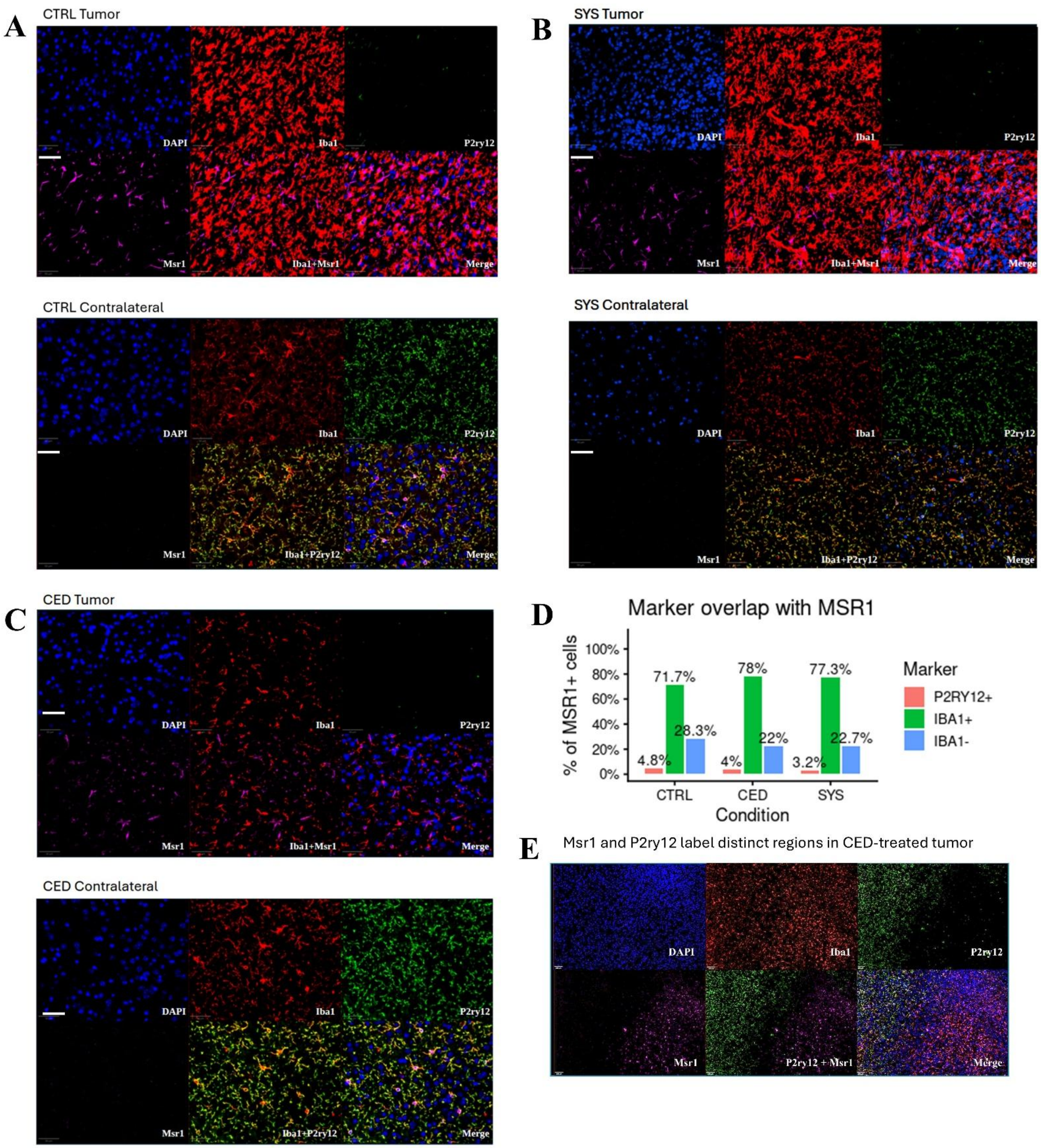


**Supplementary Figure 4: A-C** – Representative immunofluorescence images of Iba1, Msr1, and P2ry12 in the tumor and contralateral hemisphere of APCL tumor-bearing mice under control, systemic dexamethasone (10 mg/kg/day i.p.), CED-dexamethasone (100 ng/µl in 1007D Alzet pump) treatment conditions, respectively. White bar represents 50 µm. **D** – Quantification from one representative sample per condition showed that Iba1 labeling was present in the majority of Msr1+ cells, supporting their myeloid identity. In contrast, P2ry12 labeling among Msr1+ cells was rare, consistent with these markers identifying largely distinct populations. **E** – Representative image from a CED-dexamethasone-treated mouse demonstrates that Msr1 and P2ry12 label spatially distinct regions, with Msr1 concentrated in the tumor core and P2ry12 localized outside the tumor. White bar represents 100 µm.

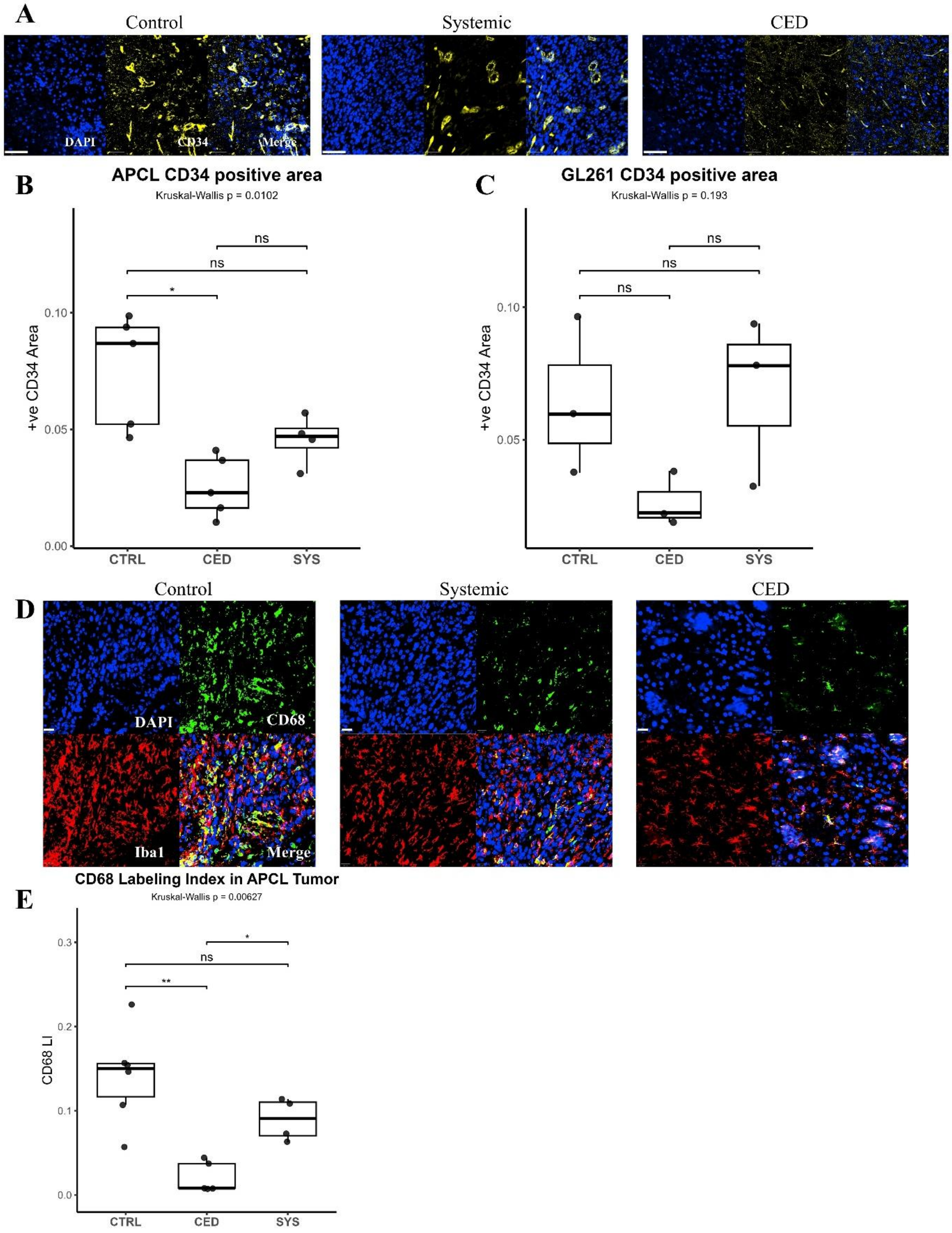


**Supplementary Figure 5**: **A** – Representative immunofluorescence images showing CD68 and CD34 in the tumor core of GL261 tumor-bearing mice. White bar represents 20 µm. **B** – Both systemic and CED dexamethasone led to a significant reduction in CD68-positive cells compared with control (CTRL vs SYS, $p = 0.0139$; CTRL vs CED, $p = 0.0228$), with no significant difference between the systemic and CED groups (SYS vs CED, $p = 0.658$). **C** – Both systemic and CED dexamethasone also led to a significant reduction in CD34-positive staining area normalized to cellularity compared with control (CTRL vs SYS, $p = 0.0134$; CTRL vs CED, $p = 0.0143$), with no significant difference between the systemic and CED groups (SYS vs CED, $p = 0.527$). Statistical testing was performed using a Kruskal–Wallis test with post-hoc pairwise Welch's t-tests (* = $p < 0.05$; ** = $p < 0.01$, *** = $p < 0.001$, **** = $p < 0.0001$).

**A**

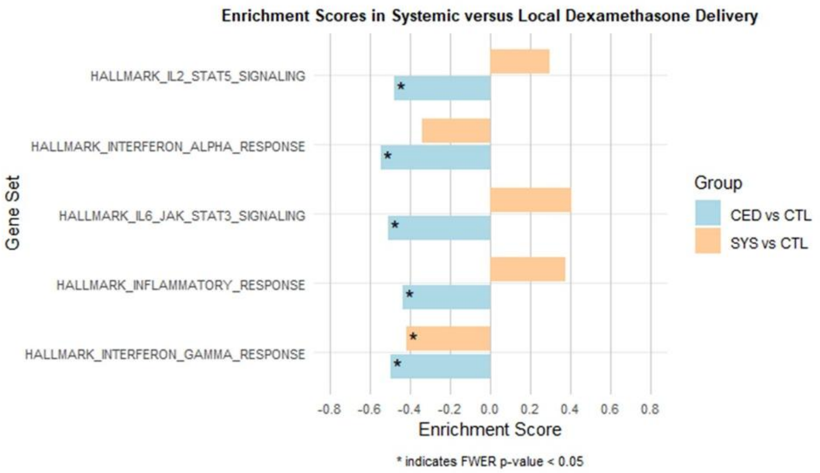


**B** Programs in Mouse Glioma Myeloid Cells Treated with CTRL vs CED and CTL vs SYS Dexamethasone

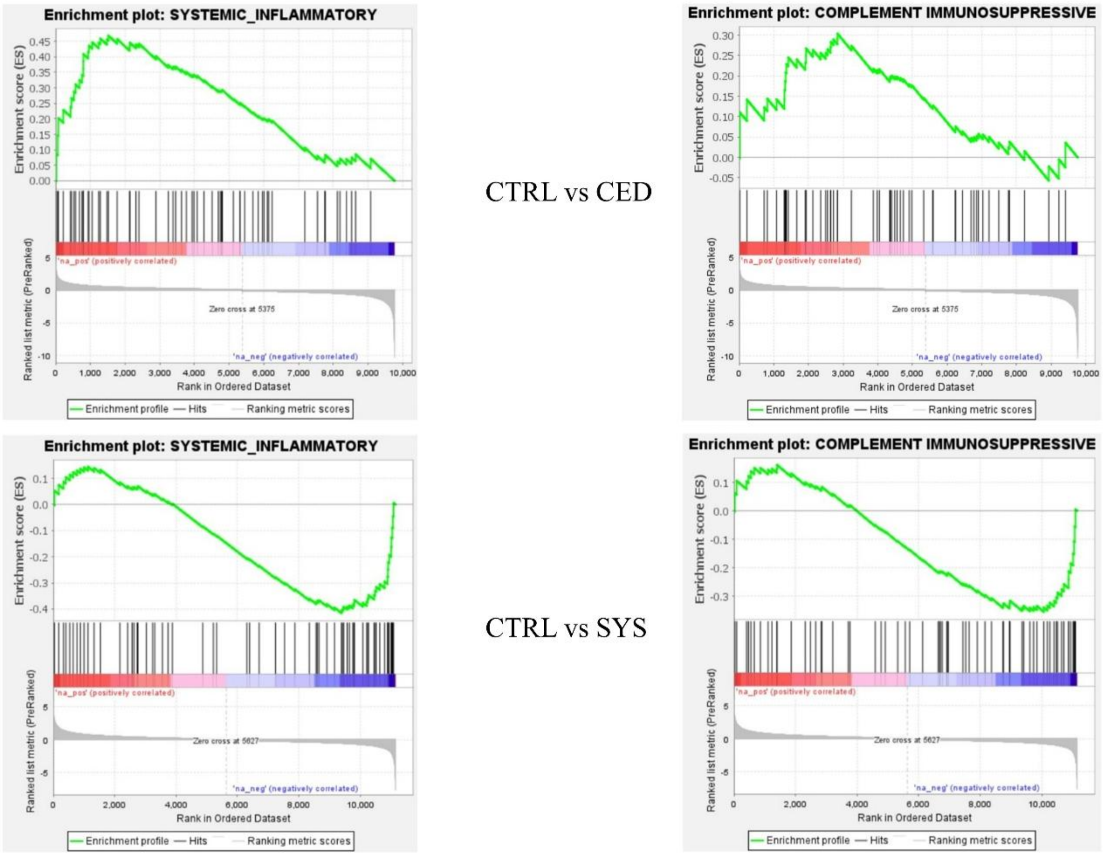


**Supplementary Figure 6**: **A** – GSEA enrichment scores for select Hallmark inflammatory pathways. Interferon response pathways moved in the same direction, regardless of delivery method, while other inflammatory gene sets moved in opposite directions. Local delivery appears to be more uniformly anti-inflammatory than once daily systemic delivery. **B** – The "complement immunosuppressive" and "systemic inflammatory" pathways from Miller et al. were differentially regulated in the local vs systemic delivery groups. Local delivery led to a NES of -1.39 whereas systemic delivery had a NES of +1.21 in the "complement immunosuppressive" gene set. Similarly, local delivery led to a NES of -1.59 whereas systemic delivery had a NES of +1.91 in the "systemic inflammatory" gene set.c

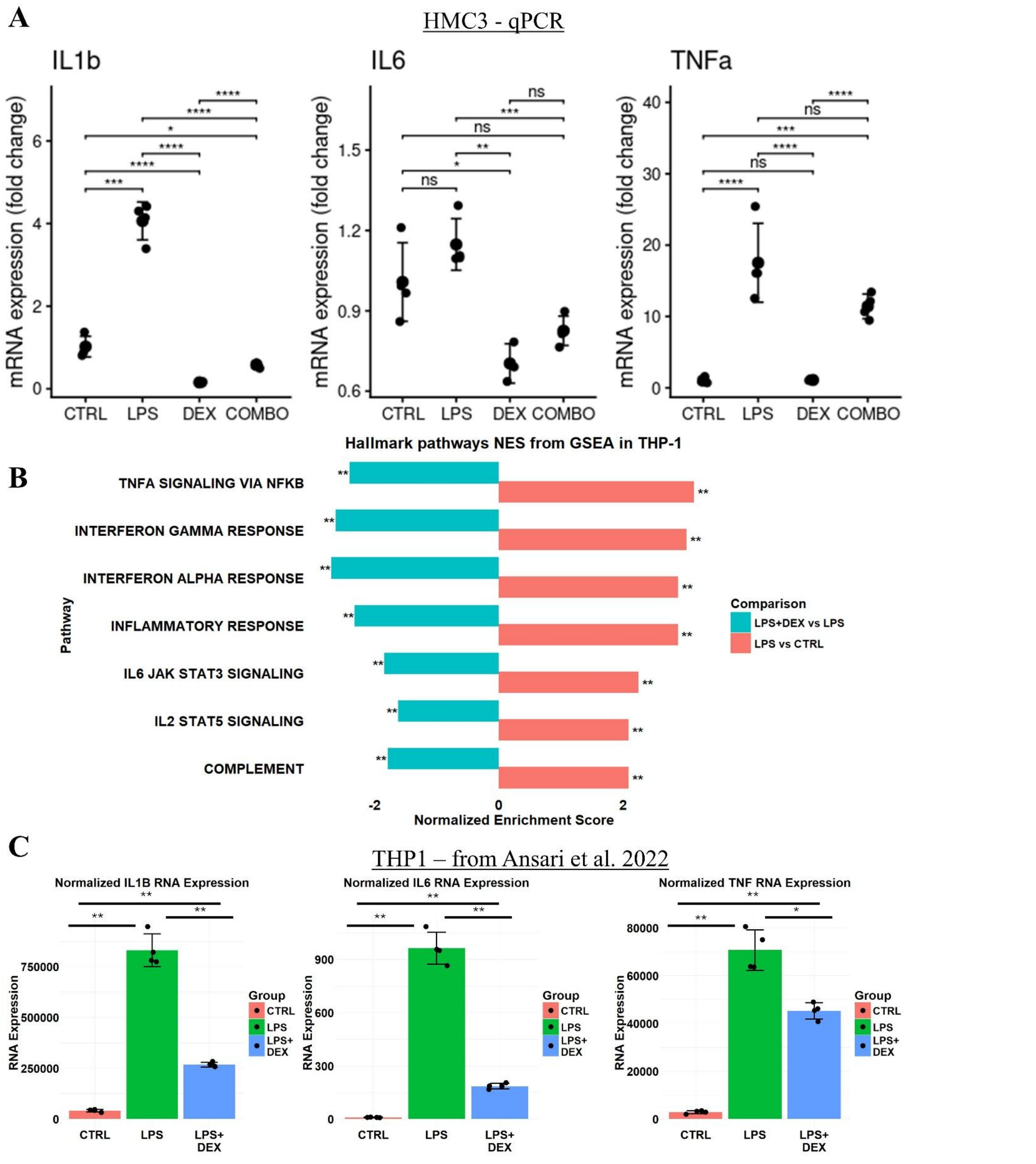


**Supplementary Figure 7**: **A** – qPCR for *IL1b, IL6,* and *TNFa* expression in human HMC3 microglia treated with vehicle, LPS, dexamethasone, or LPS+dexamethasone. **B** – LPS induced a broad upregulation of Hallmark inflammatory genes in THP-1 cells, which were relatively downregulated when cells were treated with a combination of LPS+Dex. **C** – Normalized RNA expression for IL-1β, IL6, and TNFα. Statistical testing was performed using a Kruskal–Wallis test with post-hoc pairwise Welch's t-tests (* = $p < 0.05$; ** = $p < 0.01$, *** = $p < 0.001$, **** = $p < 0.0001$)

**Supplementary Table 1**

Nominal dexamethasone concentration was 100 ng/µl.

| Days at 37C | Dexamethasone concentration (ng/µl) |
|---|---|
| 1 | 93.99 |
| 3 | 91.02 |
| 4 | 91.05 |
| 5 | 91.11 |
| 6 | 95.76 |
| 7 | 92.42 |
| 8 | 92.21 |
| 10 | 94.42 |

**Supplementary Table 1**: Dexamethasone remains stable at physiologic conditions for up to 10 days.

**Supplementary Table 2**

| Gene Name |
|---|
| ABHD12 |
| AKAP13 |
| APBB1IP |
| ASAP1 |
| BASP1 |
| BIN2 |
| CTSS |
| CD2 |
| CD74 |
| CMSS1 |
| C1QC |
| CX3CR1 |
| CST3 |
| DOCK8 |
| DAPP1 |
| FGD2 |
| GPR34 |
| GZMA |
| HPGDS |
| ITGA6 |
| IFI27 |
| IL2RA |
| LAIR1 |
| LY86 |
| MALAT1 |
| MEF2C |
| NAV2 |
| NAV3 |
| NR3C1 |
| PTN |
| PLXDC2 |
| PENK |
| RAD51B |
| ARHGAP45 |
| SCIN |
| SLAMF7 |
| SUSD3 |
| TSPAN8 |
| TREM2 |
| UNC93B1 |

**Supplementary Table 2**: Top downregulated genes in CED-treated, myeloid nuclei in the snRNA data compared to control. This gene set was used to assess the prognostic importance of suppressing an inflammatory transcriptional signature by correlating survival to the expression of these genes in the TCGA and CGGA datasets.

### Supplementary Table 3

| Gene Rank | Gene | avg_log2FC | p_val_adj |
|---|---|---|---|
| 1 | Fkbp5 | -1.458 | 1.71E-72 |
| 2 | Mir99ahg | 1.925 | 5.98E-52 |
| 3 | Ccnd3 | -1.371 | 5.58E-51 |
| 4 | Nav2 | 2.094 | 1.70E-47 |
| 5 | Slc9a9 | 1.267 | 2.25E-37 |
| 6 | Zbtb16 | -1.815 | 6.84E-37 |
| 7 | Gcnt2 | -1.985 | 1.50E-31 |
| 8 | Tanc2 | 0.957 | 2.17E-31 |
| 9 | Arl15 | -1.208 | 3.57E-31 |
| 10 | Plxdc2 | 1.205 | 9.57E-31 |
| 11 | Tmcc3 | 0.918 | 2.77E-29 |
| 12 | Cpeb2 | -1.230 | 3.18E-23 |
| 13 | Csmd3 | 1.839 | 1.31E-22 |
| 14 | Fn1 | -3.367 | 5.00E-22 |
| 15 | Rnf169 | -0.829 | 9.45E-22 |
| 16 | Klf13 | -1.074 | 1.14E-20 |
| 17 | Ap1s2 | -1.486 | 3.06E-20 |
| 18 | Lrmda | 0.938 | 1.73E-19 |
| 19 | Ctsl | -1.425 | 3.79E-18 |
| 20 | Slc24a3 | -1.213 | 7.69E-18 |
| 21 | Il4ra | -1.929 | 1.07E-17 |
| 22 | C5ar1 | -2.176 | 1.39E-17 |
| 23 | Inpp4b | 0.867 | 5.36E-17 |
| 24 | Ssh2 | -0.560 | 7.43E-17 |
| 25 | Rab27a | -1.426 | 2.03E-16 |
| 26 | Kdm7a | -0.997 | 8.61E-16 |
| 27 | Arhgap45 | -0.712 | 1.29E-15 |
| 28 | Plxna4 | 1.460 | 1.94E-15 |
| 29 | Lifr | -1.130 | 2.09E-15 |
| 30 | Il1r2 | -5.533 | 2.44E-15 |
| 31 | Pik3ip1 | -2.348 | 2.56E-15 |
| 32 | Mcl1 | -1.267 | 1.49E-14 |
| 33 | Fchsd2 | 1.180 | 1.88E-14 |
| 34 | Chil3 | -5.383 | 6.98E-14 |
| 35 | Cebpb | -1.972 | 7.65E-14 |
| 36 | Slc38a2 | -1.035 | 1.30E-13 |
| 37 | Snx29 | 0.868 | 1.82E-13 |
| 38 | Large1 | 1.184 | 2.87E-13 |
| 39 | Rnf180 | 1.416 | 3.88E-13 |
| 40 | Cd300lf | -1.376 | 4.27E-13 |
| 41 | Peli2 | -0.846 | 7.77E-13 |
| 42 | Spp1 | -3.740 | 1.24E-12 |
| 43 | Fcgr3 | -1.288 | 1.25E-12 |
| 44 | Igf1r | -0.999 | 3.62E-12 |
| 45 | Ptpn1 | -0.863 | 4.42E-12 |
| 46 | Cd244a | -1.378 | 1.04E-11 |
| 47 | Picalm | -0.566 | 1.38E-11 |
| 48 | Arhgap24 | -0.882 | 4.38E-11 |
| 49 | Ctss | -0.678 | 4.75E-11 |
| 50 | Slc39a14 | -2.048 | 5.70E-11 |
| 51 | Mir142hg | -1.058 | 6.40E-11 |
| 52 | Cept1 | -0.854 | 2.23E-10 |
| 53 | Alox5ap | -1.734 | 3.09E-10 |
| 54 | Tpd52 | -0.936 | 3.14E-10 |
| 55 | Rin3 | -0.861 | 3.72E-10 |
| 56 | Frmd4a | 0.841 | 3.74E-10 |
| 57 | Socs3 | -3.058 | 3.99E-10 |
| 58 | Aoah | 2.036 | 4.11E-10 |
| 59 | Ms4a6c | -1.392 | 5.23E-10 |
| 60 | Per1 | -2.289 | 8.56E-10 |
| 61 | Maml3 | -0.649 | 1.28E-09 |
| 62 | Herc4 | -0.760 | 2.11E-09 |
| 63 | Mafb | -0.986 | 2.17E-09 |
| 64 | Osbpl9 | -0.939 | 2.23E-09 |
| 65 | Mmp8 | -4.979 | 2.25E-09 |
| 66 | Gm13710 | 3.122 | 2.58E-09 |
| 67 | Hp | -3.319 | 3.13E-09 |
| 68 | Gm10790 | 1.147 | 3.34E-09 |
| 69 | C030034L19Rik | -1.619 | 3.34E-09 |
| 70 | Slc8a1 | 0.464 | 3.58E-09 |
| 71 | Fcer1g | -1.167 | 3.79E-09 |
| 72 | Cdk8 | -0.739 | 4.02E-09 |
| 73 | Gsr | -1.386 | 4.81E-09 |
| 74 | Tmem154 | -2.193 | 5.45E-09 |
| 75 | Tmem87b | -0.734 | 7.64E-09 |
| 76 | Notch4 | -1.699 | 7.88E-09 |
| 77 | Sgip1 | -1.113 | 9.68E-09 |
| 78 | Ripor2 | -0.983 | 9.93E-09 |
| 79 | Tsc22d3 | -0.800 | 1.60E-08 |
| 80 | Gda | -2.186 | 1.96E-08 |
| 81 | Ucp2 | -1.168 | 2.85E-08 |
| 82 | Arrb2 | -0.765 | 3.35E-08 |
| 83 | Dhx9 | -0.744 | 3.45E-08 |
| 84 | Ddi2 | -0.669 | 4.40E-08 |
| 85 | Ms4a8a | -3.338 | 5.45E-08 |
| 86 | Camk1d | -0.394 | 6.19E-08 |
| 87 | Srgap2 | 0.516 | 7.29E-08 |
| 88 | Stab1 | -0.709 | 7.99E-08 |
| 89 | Ctsd | -0.694 | 1.00E-07 |
| 90 | Rassf2 | -0.701 | 1.01E-07 |
| 91 | Cd180 | 1.433 | 1.20E-07 |
| 92 | Man2a1 | -0.627 | 1.35E-07 |
| 93 | Nfam1 | -0.891 | 1.42E-07 |
| 94 | F10 | -3.302 | 1.50E-07 |
| 95 | Inf2 | -1.580 | 1.74E-07 |
| 96 | Ston1 | -1.567 | 1.78E-07 |
| 97 | Ms4a4a | -1.544 | 1.80E-07 |
| 98 | Fxyd5 | -1.902 | 1.91E-07 |
| 99 | Rcsd1 | -0.710 | 1.96E-07 |
| 100 | Sla | -0.693 | 2.02E-07 |
| 101 | Igsf6 | -1.179 | 2.46E-07 |
| 102 | Ms4a6d | -1.102 | 2.67E-07 |
| 103 | Clk4 | -0.897 | 2.69E-07 |
| 104 | Srgn | -2.706 | 2.78E-07 |
| 105 | Cmss1 | -0.595 | 2.94E-07 |
| 106 | Ccr2 | -2.101 | 3.08E-07 |
| 107 | Ccdc192 | 3.420 | 3.52E-07 |
| 108 | Snx1 | -0.901 | 4.01E-07 |
| 109 | Bin2 | -0.650 | 4.78E-07 |
| 110 | Sbno2 | -0.871 | 6.93E-07 |
| 111 | Arpc1b | -1.024 | 8.06E-07 |
| 112 | Frrs1 | -0.746 | 8.39E-07 |
| 113 | S100a10 | -2.911 | 9.63E-07 |
| 114 | Cast | -1.265 | 9.81E-07 |
| 115 | Pde2a | -0.873 | 1.06E-06 |
| 116 | Pag1 | -0.463 | 1.13E-06 |
| 117 | Emilin2 | -2.439 | 1.23E-06 |
| 118 | Ctsa | -0.920 | 1.25E-06 |
| 119 | Cep152 | -1.017 | 1.27E-06 |
| 120 | Mmp19 | -4.773 | 1.30E-06 |
| 121 | Serp1 | -1.365 | 1.40E-06 |
| 122 | Hnrnpf | -0.905 | 1.41E-06 |
| 123 | Il17ra | -1.070 | 1.49E-06 |
| 124 | Gphn | -0.582 | 1.62E-06 |
| 125 | Rbm47 | -0.374 | 1.63E-06 |
| 126 | Slc43a2 | -0.640 | 1.63E-06 |
| 127 | Fcgr2b | -0.960 | 1.67E-06 |
| 128 | H2-D1 | -0.914 | 1.91E-06 |
| 129 | Tcn2 | -1.216 | 2.34E-06 |
| 130 | Nufip2 | -0.582 | 2.70E-06 |
| 131 | Ttc28 | 0.881 | 2.71E-06 |
| 132 | Cux1 | -0.519 | 2.91E-06 |
| 133 | Srsf7 | -0.705 | 3.03E-06 |
| 134 | Ophn1 | 0.556 | 3.10E-06 |
| 135 | Washc2 | -0.805 | 3.15E-06 |
| 136 | Tgfbi | -2.264 | 3.26E-06 |
| 137 | Mia2 | -0.620 | 3.38E-06 |
| 138 | Ly86 | 0.759 | 3.46E-06 |
| 139 | Atrnl1 | -1.001 | 3.81E-06 |
| 140 | Mafg | -0.726 | 3.92E-06 |
| 141 | Comt | -0.513 | 4.47E-06 |
| 142 | Diaph1 | -0.993 | 4.54E-06 |
| 143 | Ogfrl1 | -0.706 | 5.93E-06 |
| 144 | Mt2 | -1.067 | 5.97E-06 |
| 145 | Tbc1d15 | -1.084 | 6.01E-06 |
| 146 | Itga9 | 1.430 | 7.18E-06 |
| 147 | Polg | -0.856 | 7.25E-06 |
| 148 | Hmox1 | -1.613 | 8.18E-06 |
| 149 | Rbm26 | -0.564 | 8.25E-06 |
| 150 | Clec4d | -3.201 | 8.28E-06 |
| 151 | Cadm1 | 0.950 | 8.28E-06 |
| 152 | Gm19951 | -0.770 | 8.46E-06 |
| 153 | Tet2 | -0.637 | 8.82E-06 |
| 154 | Qki | 0.440 | 9.29E-06 |
| 155 | Swt1 | -0.837 | 9.42E-06 |
| 156 | Nfkbia | -1.240 | 9.51E-06 |
| 157 | Tg | -2.710 | 9.63E-06 |
| 158 | Ccr1 | -2.508 | 1.01E-05 |
| 159 | Hdac4 | -0.897 | 1.05E-05 |
| 160 | Dok3 | -1.331 | 1.06E-05 |
| 161 | Fndc3b | -0.642 | 1.13E-05 |
| 162 | Il15ra | -0.956 | 1.14E-05 |
| 163 | Stat3 | -0.684 | 1.15E-05 |
| 164 | Gm56663 | -4.042 | 1.23E-05 |
| 165 | Vsir | -0.501 | 1.24E-05 |
| 166 | Meg3 | 0.756 | 1.31E-05 |
| 167 | Ptprm | 1.180 | 1.44E-05 |
| 168 | Zdhhc9 | -1.471 | 1.63E-05 |
| 169 | 2210408F21Rik | -0.890 | 2.11E-05 |
| 170 | Klhl24 | -0.909 | 2.12E-05 |
| 171 | Thbs1 | -4.183 | 2.19E-05 |
| 172 | Fbxl20 | -0.565 | 2.21E-05 |
| 173 | Filip1l | -0.531 | 2.22E-05 |
| 174 | Jdp2 | -1.082 | 2.29E-05 |
| 175 | Pgd | -1.096 | 2.31E-05 |
| 176 | Msrb1 | -2.143 | 2.75E-05 |
| 177 | Btg1 | -1.136 | 2.91E-05 |
| 178 | Ifitm3 | -2.200 | 2.99E-05 |
| 179 | Otub1 | -1.988 | 3.05E-05 |
| 180 | Lpcat2 | 0.796 | 3.18E-05 |
| 181 | Tm6sf1 | -0.426 | 3.22E-05 |
| 182 | Zfp697 | -0.958 | 3.29E-05 |
| 183 | Tyrobp | -1.616 | 3.96E-05 |
| 184 | Ltb4r1 | -4.649 | 3.97E-05 |
| 185 | Abhd15 | -0.786 | 4.00E-05 |
| 186 | Parp4 | -0.982 | 4.06E-05 |
| 187 | Calr | -0.981 | 4.31E-05 |
| 188 | Ddit4 | -1.367 | 4.83E-05 |
| 189 | Man2b1 | -0.625 | 5.01E-05 |
| 190 | Map3k8 | -0.896 | 5.12E-05 |
| 191 | Ifngr1 | -0.527 | 5.36E-05 |
| 192 | Dmxl2 | -0.927 | 6.06E-05 |
| 193 | Zfp36l2 | -0.826 | 6.06E-05 |
| 194 | Osbpl11 | -0.528 | 6.43E-05 |
| 195 | Svil | -0.919 | 6.46E-05 |
| 196 | Npnt | 2.890 | 6.90E-05 |
| 197 | Tpp2 | -0.678 | 6.92E-05 |
| 198 | Klhl2 | -0.931 | 6.93E-05 |
| 199 | Adgre1 | 1.278 | 6.99E-05 |
| 200 | Dusp3 | -0.847 | 7.26E-05 |
| 201 | Cdk14 | -0.505 | 7.61E-05 |
| 202 | Kcnma1 | 1.286 | 8.05E-05 |
| 203 | Neat1 | -1.254 | 8.33E-05 |
| 204 | Dram2 | -0.462 | 8.71E-05 |
| 205 | Sell | -4.102 | 1.07E-04 |
| 206 | Atp8b4 | -1.301 | 1.14E-04 |
| 207 | Pkn1 | -0.557 | 1.29E-04 |
| 208 | Jak1 | -0.496 | 1.32E-04 |
| 209 | Klf9 | -0.966 | 1.35E-04 |
| 210 | Pfkfb3 | -0.707 | 1.37E-04 |
| 211 | Tspan5 | -0.694 | 1.38E-04 |
| 212 | Wdr26 | -0.487 | 1.41E-04 |
| 213 | Chd9 | 0.496 | 1.55E-04 |
| 214 | S100a11 | -3.531 | 1.61E-04 |
| 215 | Knop1 | -0.751 | 1.67E-04 |
| 216 | Hif3a | -1.886 | 1.76E-04 |
| 217 | Vcan | -2.863 | 1.83E-04 |
| 218 | Ppp1r10 | -0.677 | 1.84E-04 |
| 219 | Tmod1 | -1.093 | 1.90E-04 |
| 220 | Stat1 | 1.301 | 2.09E-04 |
| 221 | Elmo1 | 0.272 | 2.11E-04 |
| 222 | Il6ra | -0.540 | 2.18E-04 |
| 223 | Rnf149 | -0.817 | 2.31E-04 |
| 224 | H2-K1 | -0.738 | 2.45E-04 |
| 225 | Sirpa | 0.665 | 2.49E-04 |
| 226 | Aopep | -0.558 | 2.85E-04 |
| 227 | Iigp1 | 2.981 | 2.87E-04 |
| 228 | Hspa5 | -0.683 | 3.00E-04 |
| 229 | Slc30a5 | -0.926 | 3.17E-04 |
| 230 | Grk6 | -1.174 | 3.60E-04 |
| 231 | Slc2a1 | -1.582 | 3.76E-04 |
| 232 | Shisa5 | -1.500 | 4.24E-04 |
| 233 | Stx5a | -0.674 | 4.35E-04 |
| 234 | Mt1 | -0.614 | 4.45E-04 |
| 235 | Smpdl3a | -1.882 | 5.07E-04 |
| 236 | Pkm | -0.582 | 5.10E-04 |
| 237 | Gpi1 | -0.619 | 5.25E-04 |
| 238 | Grn | -0.691 | 5.39E-04 |
| 239 | Sdc4 | -1.265 | 5.69E-04 |
| 240 | Esco1 | -0.568 | 5.79E-04 |
| 241 | Zbed6 | -0.852 | 5.81E-04 |
| 242 | Ube2h | -0.458 | 5.91E-04 |
| 243 | Atp8a1 | -0.464 | 5.93E-04 |
| 244 | Slc2a3 | -0.601 | 6.08E-04 |
| 245 | Csf2rb | -0.854 | 6.16E-04 |
| 246 | Sort1 | -0.489 | 6.19E-04 |
| 247 | Tenm2 | 1.830 | 6.58E-04 |
| 248 | Hipk1 | -0.756 | 6.70E-04 |
| 249 | S100a6 | -2.519 | 6.76E-04 |
| 250 | Alpk1 | -1.089 | 6.78E-04 |
| 251 | Sipa1 | -0.398 | 7.02E-04 |
| 252 | Ldlrad4 | 0.490 | 7.08E-04 |
| 253 | 1600014C10Rik | -1.053 | 7.14E-04 |
| 254 | Rassf3 | -0.556 | 7.85E-04 |
| 255 | Arg1 | -3.250 | 8.33E-04 |
| 256 | Pdk1 | -0.476 | 8.52E-04 |
| 257 | Nisch | -0.516 | 9.03E-04 |
| 258 | Hbp1 | -0.609 | 9.09E-04 |
| 259 | Xbp1 | -0.998 | 9.32E-04 |
| 260 | Gpr146 | -1.018 | 9.65E-04 |
| 261 | Lrp1b | 1.869 | 9.74E-04 |
| 262 | Prkag2 | -0.776 | 9.89E-04 |
| 263 | Apoc2 | -3.869 | 1.03E-03 |
| 264 | Slfn2 | -1.161 | 1.10E-03 |
| 265 | Rc3h1 | -0.576 | 1.14E-03 |
| 266 | Fbxl5 | -0.850 | 1.15E-03 |
| 267 | Itgb2 | -0.662 | 1.17E-03 |
| 268 | Plet1os | -2.536 | 1.20E-03 |
| 269 | Rnpep | -0.640 | 1.25E-03 |
| 270 | Cd53 | -0.712 | 1.27E-03 |

| | | | |
|---|---|---|---|
| 271 | Apobec1 | -0.415 | 1.28E-03 |
| 272 | Tmem156 | -1.363 | 1.34E-03 |
| 273 | Mast3 | -0.700 | 1.36E-03 |
| 274 | Npepps | -0.667 | 1.37E-03 |
| 275 | Hacd2 | -0.731 | 1.37E-03 |
| 276 | Scarb1 | -0.870 | 1.38E-03 |
| 277 | Gm33100 | -1.164 | 1.41E-03 |
| 278 | Atrn | -0.859 | 1.49E-03 |
| 279 | Dpp8 | -0.595 | 1.53E-03 |
| 280 | Trappc14 | -0.586 | 1.54E-03 |
| 281 | Malat1 | 0.129 | 1.67E-03 |
| 282 | Pygl | -1.624 | 1.68E-03 |
| 283 | Ostf1 | -0.543 | 1.68E-03 |
| 284 | Tiparp | -1.532 | 1.73E-03 |
| 285 | Clk1 | -0.440 | 1.77E-03 |
| 286 | Pirb | -1.519 | 1.79E-03 |
| 287 | Fth1 | 0.136 | 1.79E-03 |
| 288 | Lars2 | -0.562 | 1.81E-03 |
| 289 | Chaserr | -0.494 | 1.82E-03 |
| 290 | Rbm5 | -0.434 | 1.84E-03 |
| 291 | Prelid1 | -1.556 | 1.87E-03 |
| 292 | Emb | -1.778 | 1.88E-03 |
| 293 | Eed | -0.637 | 1.91E-03 |
| 294 | Ctsz | -0.526 | 1.94E-03 |
| 295 | Ibtk | -1.320 | 1.96E-03 |
| 296 | Ctsh | -0.477 | 1.99E-03 |
| 297 | Maml2 | 0.548 | 2.02E-03 |
| 298 | Cyba | -1.058 | 2.10E-03 |
| 299 | Macir | -0.929 | 2.13E-03 |
| 300 | B4galt1 | -0.365 | 2.14E-03 |
| 301 | Tmbim6 | -0.396 | 2.16E-03 |
| 302 | Baz1a | -0.417 | 2.26E-03 |
| 303 | Stk24 | -0.598 | 2.30E-03 |
| 304 | Plaur | -1.409 | 2.35E-03 |
| 305 | Arhgef18 | -0.905 | 2.42E-03 |
| 306 | Zfc3h1 | -0.461 | 2.46E-03 |
| 307 | Ahnak | -1.426 | 2.48E-03 |
| 308 | Wipf3 | -1.339 | 2.50E-03 |
| 309 | Cd63 | -0.997 | 2.51E-03 |
| 310 | Nrxn3 | 0.653 | 2.61E-03 |
| 311 | Snx18 | -0.709 | 2.67E-03 |
| 312 | Rab21 | -0.547 | 2.77E-03 |
| 313 | Sorl1 | -0.551 | 2.82E-03 |
| 314 | Pak2 | -0.417 | 2.87E-03 |
| 315 | Tns3 | 0.658 | 2.89E-03 |
| 316 | Cd68 | -1.099 | 2.93E-03 |
| 317 | Il18rap | -2.097 | 2.97E-03 |
| 318 | Arhgap30 | -0.513 | 3.01E-03 |
| 319 | Selplg | -0.478 | 3.33E-03 |
| 320 | Sirpb1c | -2.454 | 3.39E-03 |
| 321 | Capg | -1.971 | 3.56E-03 |
| 322 | Gnai2 | -0.750 | 3.73E-03 |
| 323 | Smad4 | -0.511 | 3.81E-03 |
| 324 | Irak4 | -0.652 | 3.82E-03 |
| 325 | Dock2 | 0.432 | 3.87E-03 |
| 326 | Iqgap1 | -0.779 | 4.00E-03 |
| 327 | Prpf40a | -0.585 | 4.04E-03 |
| 328 | Csad | -0.759 | 4.21E-03 |
| 329 | Laptm5 | -0.525 | 4.39E-03 |
| 330 | Cebpg | -0.569 | 4.40E-03 |
| 331 | Elf4 | -0.461 | 4.56E-03 |
| 332 | Tcp11l2 | -1.014 | 4.68E-03 |
| 333 | Mau2 | -0.459 | 4.85E-03 |
| 334 | Arpc2 | -0.396 | 5.03E-03 |
| 335 | Plod1 | -0.616 | 5.07E-03 |
| 336 | Rsrp1 | -0.437 | 5.14E-03 |
| 337 | 4930447J18Rik | -4.951 | 5.16E-03 |
| 338 | Coro1a | -0.854 | 5.19E-03 |
| 339 | Napsa | -3.087 | 5.34E-03 |
| 340 | Mlph | -0.629 | 5.39E-03 |
| 341 | Adamtsl4 | -1.598 | 5.53E-03 |
| 342 | Fmnl1 | -0.630 | 5.55E-03 |
| 343 | Rnf145 | -0.674 | 5.59E-03 |
| 344 | Coro2a | -0.482 | 5.77E-03 |
| 345 | Dapp1 | -0.412 | 5.80E-03 |
| 346 | Slc16a6 | -0.651 | 5.86E-03 |
| 347 | Nfil3 | -2.408 | 6.08E-03 |
| 348 | Vgll4 | -0.538 | 6.17E-03 |
| 349 | Gm48633 | -7.700 | 6.21E-03 |
| 350 | Ms4a4c | -1.720 | 6.28E-03 |
| 351 | Kmt2e | -0.332 | 6.50E-03 |
| 352 | Txnip | -0.959 | 6.52E-03 |
| 353 | Zfp740 | -0.636 | 6.54E-03 |
| 354 | Fhit | 1.188 | 6.65E-03 |
| 355 | Flot1 | -1.242 | 6.70E-03 |
| 356 | Rab5if | -0.982 | 7.16E-03 |
| 357 | Snrnp70 | -0.380 | 7.29E-03 |
| 358 | Rpgrip1 | -0.916 | 7.31E-03 |
| 359 | Ifi209 | -1.350 | 7.59E-03 |
| 360 | Cebpd | -1.447 | 7.69E-03 |
| 361 | Pdia3 | -0.434 | 7.84E-03 |
| 362 | Abcg1 | -0.487 | 7.91E-03 |
| 363 | Epb41l2 | 0.495 | 8.11E-03 |
| 364 | Rab32 | -0.999 | 8.29E-03 |
| 365 | Tars2 | -1.213 | 8.49E-03 |
| 366 | Nipa2 | -0.563 | 8.52E-03 |
| 367 | Abca6 | -3.576 | 8.67E-03 |
| 368 | Sell1 | -0.521 | 8.69E-03 |
| 369 | Stk17b | -0.755 | 8.80E-03 |
| 370 | Pnkp | -1.741 | 8.81E-03 |
| 371 | C1qa | -0.591 | 9.08E-03 |
| 372 | Fes | -0.825 | 9.26E-03 |
| 373 | Bst1 | -2.790 | 9.32E-03 |
| 374 | Agl | -0.747 | 9.41E-03 |
| 375 | Setx | -0.703 | 9.42E-03 |
| 376 | Aff1 | -0.447 | 9.51E-03 |
| 377 | Gm46224 | -0.607 | 9.94E-03 |
| 378 | Gcnt1 | -0.494 | 9.94E-03 |
| 379 | Tent5a | -0.520 | 1.02E-02 |
| 380 | Ecm1 | -1.752 | 1.06E-02 |
| 381 | Trf | -0.419 | 1.07E-02 |
| 382 | Ms4a6b | -0.430 | 1.10E-02 |
| 383 | Faf2 | -0.853 | 1.12E-02 |
| 384 | Lrp10 | -0.859 | 1.14E-02 |
| 385 | Ppard | -0.637 | 1.15E-02 |
| 386 | Ypel3 | -0.412 | 1.16E-02 |
| 387 | Anxa2 | -2.047 | 1.16E-02 |
| 388 | Prdm2 | -0.492 | 1.19E-02 |
| 389 | Mlxip | -0.389 | 1.19E-02 |
| 390 | Tut7 | -0.418 | 1.19E-02 |
| v391 | Phf3 | -0.384 | 1.27E-02 |
| 392 | Sult1a1 | -0.629 | 1.31E-02 |
| 393 | U2af1 | -0.727 | 1.39E-02 |
| 394 | Uhrf1bp1l | -0.736 | 1.40E-02 |
| 395 | Eif4ebp2 | -0.655 | 1.45E-02 |
| 396 | Tspan18 | 2.603 | 1.64E-02 |
| 397 | Ly6c2 | -3.861 | 1.65E-02 |
| 398 | Wfdc17 | -2.947 | 1.69E-02 |
| 399 | Bclaf1 | -0.340 | 1.73E-02 |
| 400 | Lrpap1 | -0.677 | 1.81E-02 |
| 401 | Rmdn1 | -0.510 | 1.82E-02 |
| 402 | Arpc4 | -0.738 | 1.83E-02 |
| 403 | Bank1 | 1.307 | 1.86E-02 |
| 404 | Ssbp1 | -0.531 | 1.89E-02 |
| 405 | Ifi30 | -1.402 | 1.91E-02 |
| 406 | Slfn1 | -2.085 | 2.04E-02 |
| 407 | Gm57206 | -0.849 | 2.05E-02 |
| 408 | Atp1a1 | -0.498 | 2.05E-02 |
| 409 | Ifitm2 | -2.759 | 2.07E-02 |
| 410 | Tgfbr1 | 0.326 | 2.16E-02 |
| 411 | Sfxn5 | -1.062 | 2.21E-02 |
| 412 | Cyrib | -0.289 | 2.22E-02 |
| 413 | Ripk2 | -0.608 | 2.25E-02 |
| 414 | Fam168a | -0.442 | 2.28E-02 |
| 415 | Hnrnpll | -0.767 | 2.29E-02 |
| 416 | Grid2 | 1.976 | 2.29E-02 |
| 417 | Syk | -0.481 | 2.30E-02 |
| 418 | Mdga2 | 2.059 | 2.30E-02 |
| 419 | Adam9 | -0.878 | 2.31E-02 |
| 420 | Calhm2 | -0.930 | 2.34E-02 |
| 421 | Ccar1 | -0.539 | 2.36E-02 |
| 422 | AB124611 | -0.940 | 2.38E-02 |
| 423 | Saa3 | 0.235 | 2.51E-02 |
| 424 | Psmd12 | -0.705 | 2.53E-02 |
| 425 | Tnfsf13 | -1.848 | 2.53E-02 |
| 426 | Uckl1 | -0.873 | 2.57E-02 |
| 427 | Adam8 | -2.150 | 2.68E-02 |
| 428 | Ptprj | 0.392 | 2.71E-02 |
| 429 | Peli1 | 0.759 | 2.72E-02 |
| 430 | Pitpna | -0.469 | 2.95E-02 |
| 431 | Pfn1 | -0.855 | 2.97E-02 |
| 432 | Patl1 | -0.695 | 2.98E-02 |
| 433 | Mon2 | -0.438 | 3.02E-02 |
| 434 | Rbms1 | -0.431 | 3.02E-02 |
| 435 | Ppan | -2.345 | 3.15E-02 |
| 436 | Arl6ip1 | -0.397 | 3.34E-02 |
| 437 | Adssl1 | -1.916 | 3.36E-02 |
| 438 | Cfh | -0.575 | 3.36E-02 |
| 439 | D530033B14Rik | -1.858 | 3.37E-02 |
| 440 | Hmg20b | -0.928 | 3.48E-02 |
| 441 | Gm22146 | -1.265 | 3.52E-02 |
| 442 | Slc38a1 | -0.361 | 3.61E-02 |
| 443 | Brd8 | -0.409 | 3.63E-02 |
| 444 | Hnrnpdl | -0.458 | 3.92E-02 |
| 445 | Serinc3 | -0.327 | 3.93E-02 |
| 446 | Ifi207 | -0.692 | 3.96E-02 |
| 447 | Sdcbp | -0.417 | 3.96E-02 |
| 448 | Scamp1 | -0.704 | 4.00E-02 |
| 449 | Nars | -0.544 | 4.00E-02 |
| 450 | Crot | -0.675 | 4.13E-02 |
| 451 | Ncf1 | -0.369 | 4.15E-02 |
| 452 | Dlg2 | 1.063 | 4.16E-02 |
| 453 | Crk | -0.545 | 4.28E-02 |
| 454 | Ero1b | -1.064 | 4.28E-02 |
| 455 | Scyl3 | -0.902 | 4.44E-02 |
| 456 | Pign | -0.405 | 4.49E-02 |
| 457 | Nbr1 | -0.376 | 4.60E-02 |
| 458 | Sp3 | -0.499 | 4.71E-02 |
| 459 | Acp5 | -3.136 | 4.82E-02 |
| 460 | 3110082I17Rik | -0.465 | 4.86E-02 |
| 461 | C1qc | -0.427 | 4.86E-02 |
| 462 | Eif5 | -0.532 | 4.92E-02 |
| 463 | Secisbp2 | -0.501 | 4.99E-02 |

**Supplementary Table 3**: Differentially expressed genes in myeloid cells between mice treated with CED-dexamethasone and systemic dexamethasone by snRNA-sequencing.
